# Changes in miRNA secondary structure can predict mutations associated with cancer and other diseases

**DOI:** 10.1101/2024.06.19.599688

**Authors:** Javor K. Novev, Sebastian E. Ahnert

## Abstract

MicroRNAs (miRNAs) are ubiquitous short RNAs regulating gene expression in many organisms, including humans. How the secondary structure (SS) of a mature miRNA affects its regulatory function remains an open question. Here we investigate this question through computational SS predictions of miRNA point mutants. We explore the mutational neighborhoods of miRNAs with association to human diseases, including cancer. We focus on possible SS changes independent of target-site complementarity, by leaving the seed region unchanged. We formulate metrics of the SS differences between such mutants and their wild types (WTs), and test whether these metrics predict disease association by comparing our results with the miRNASNP-v3 database. We find that disease-related mutants tend to have a higher probability of being fully unfolded than their WT; this and other SS-related measures are statistically significant at the database level. With the same approach, we identify a subset of individual miRNAs for which SS changes are most likely to predict disease-related mutations. These are hsa-miR-1269b, hsa-miR-4537, hsa-miR-4477b, hsa-miR-4641, and hsa-miR-6821-3p. In addition, we show that there are pairs of known miRNA WTs differing only by disease-related point mutations outside the seed region and exhibit very different SS. These pairs include hsa-miR-1269a—hsa-miR-1269b, and hsa-miR-3689a-3p—hsa-miR-3689b-3p.

## 1 Introduction

MicroRNAs (miRNAs) are short (typically 19-27 nt [1]) regulatory molecules that are highly conserved across many species [2]. They regulate gene expression by entering an RNA-induced silencing complex (RISC)[3], which typically then binds to the 3’ untranslated region (UTR) of mRNAs and inhibits their translation [4–7]; however, recent studies have also reported miRNA-induced translational activation via 5’ UTR binding [6, 8]. In humans and animals this binding canonically only occurs in the ‘seed’ region of the miRNA (nt 2-7) [9], though partial complementary binding to nucleotides outside this region is also common [10, 11], and modes of binding that do not involve the seed have also been recorded [12]. A single miRNA may therefore regulate a broad range of genes [2, 3]. Abnormal miRNA expression is observed in many diseases [4], including Parkinson’s disease [13] and cancer [5, 9, 13–15]. Moreover, individual miRNAs typically have either oncogenic or tumour-suppressive effects [3, 5], but there is evidence that miRNA expression is globally suppressed in tumour cells [4, 5]. However, it has also been reported that individual miRNAs [16] and miRNA families [17] can act as both oncogenes and tumour suppressors depending on context.

RNA molecules can fold due to pairing between some of their nucleobases, which can be described in terms of RNA secondary and tertiary structures. The role of miRNA secondary structure in the context of miRNA function has received some attention in the past, much of which has focussed on miRNA precursors, also known as primary miRNAs (pri-miRNAs) or pre-miRNAs depending on their processing stage [2, 18, 19]. In particular, Diederichs and Haber, who searched for tumor-associated mutations in cancer-derived cell lines, found no mutations within the mature miRNA sequences, and even though some of the pri-miRNA mutations dramatically altered secondary structure, they did not have an effect on *in vivo* processing and maturation [18]. Belter et al. [20] showed that some mature miRNAs form stable secondary structures and suggested that these play a functional role, and Sun et al. [21] investigated how single-nucleotide polymorphisms (SNPs) in mature miRNAs affect their function, noting that the variant miR-502-C/G produces a bulge that changes the structure of the pre-miRNA’s stem and likely affects the latter’s processing; a similar effect has also been observed in miR-125a [22]. Confoundingly, there are some examples of SNPs that are associated with increased risk of some diseases but decreased risks for others - the polymorphism rs2910164 in miR-146a-3p is known to predispose carriers to breast cancer, glioma, and gastric cancer, but at the same time is protective against prostate and gastric cancer [23].

The generally low frequency of mutations in mature miRNAs has been commented on by Bracken et al. [7], who presume this is due to their small size. SNPs, particularly ones in the seed region, are associated with many diseases [24]. In reviewing the literature on genetic variations in miRNAs, Borel and Anonarakis remark that data indicates a low density of polymorphisms in the seed region, suggesting a selective constraint [25].

To our knowledge, however, no study has systematically explored mutation-induced secondary structure changes in mature miRNA secondary structure or attempted to analyze the association between such changes and disease; it is this knowledge gap that we address in the present work. We hypothesize that miRNA activity could be influenced by SS, because folding of mature miRNA may reduce its binding affinity to target mRNAs and might influence its activity within RISC, as illustrated schematically in Figure 1.

**Figure 1:**
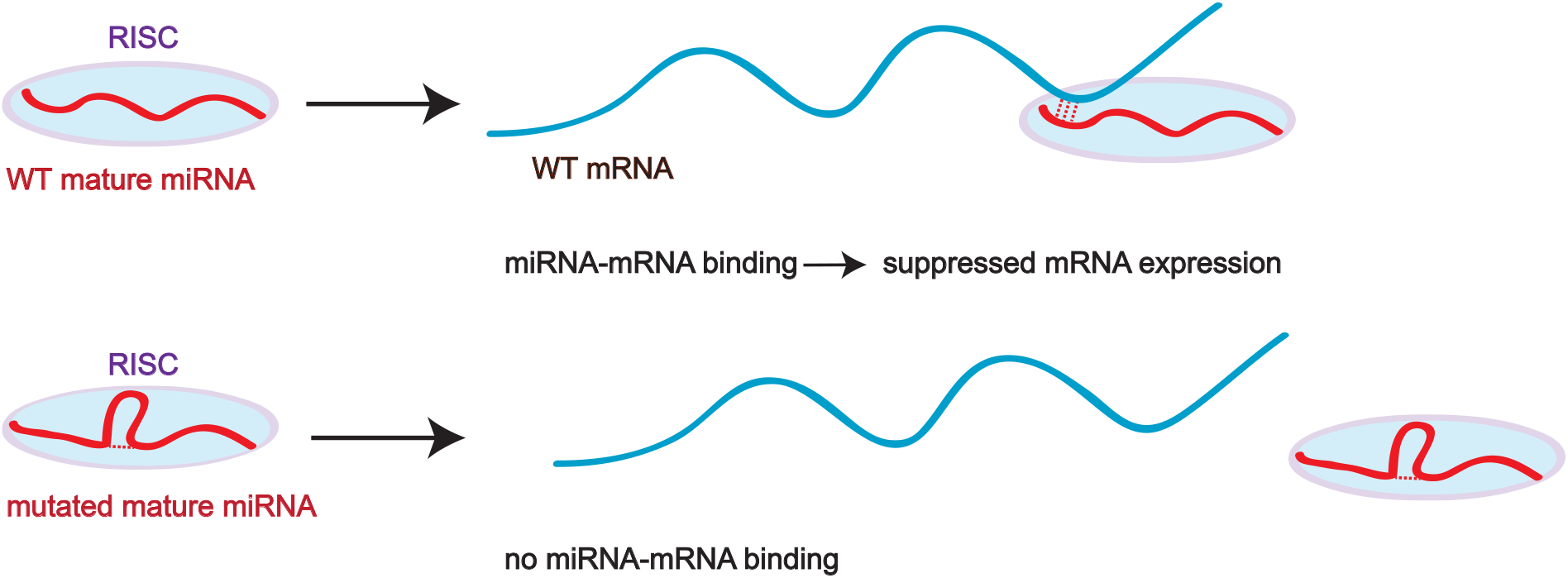
We hypothesize that miRNA mutations outside the seed region that affect miRNA secondary structure can affect miRNA-mRNA binding and thus gene expression. Illustration of the proposed mechanism by which mature miRNA SS may influence miRNA activity. Folding of the mutant miRNA (bottom) reduces its affinity to target mRNAs in comparison with the WT (top).

We study the effect of secondary structure in mature miRNAs by using computational predictions of the RNA secondary structure of point mutants for which the seed region is preserved. We then formulate multiple quantitative criteria for the changes in SS with respect to the wild type (WT). Thereafter, we test how well these criteria can predict which mutations are associated with disease. We do so by ranking the mutated sequences by a given criterion of SS change and assess whether disease-related mutants are ranked higher than expected by chance. It is not immediately clear to what extent the SS of the mature miRNA affects its activity, since the single-stranded mature miRNA is typically bound to RISC, which may affect its folding. Prior to RISC binding, a pri-miRNA is converted into a pre-miRNA by the ribonuclease III enzyme Drosha and converted into a double-stranded RNA of length 22 nt, which binds to Argonaute (Ago) proteins to form RISC [24, 26]. However, known crystal structures of RISC proteins indicate that the miRNAs within them can form base pairs [27], which suggests that mature miRNA secondary structure may play a role in miRNA activity.

While disease-associated miRNAs exhibit fewer SNPs than non-disease-related miRNAs [28], there is a substantial amount of data on the association of specific miRNA point mutations with disease. We used a database of this kind to examine the changes in secondary structure as a result of disease-related mutations. This database is miRNASNP-v3 [29], which contains data on miRNA mutations related to various diseases. We also analysed the somatic mutations in cancer collected in the SomamiR 2.0 database [30], but as almost all of them are covered in miRNASNP-v3, we only discuss these results in the Supplementary Information. Note that other online databases with mutations in miRNAs exist. For an overview see [31].

The miRNASNP-v3 database, which we use in its version from 14th May 2024, contains 2613 distinct entries recorded for the mature region. We filtered the mutations in the miRNASNP-v3 database by several criteria. Firstly we filter out the 17 entries in miRNASNP-v3 for which the related disease was unspecified, and a further 34 mutations for which the WT miRNA sequence in the miRNASNP-v3 entry is not an exact match for the one given in miRBase [32], or for which the mutated miRNA sequence specified in miRNASNP-v3 does not match the one derived from mutating the corresponding DNA sequence in the Genome Reference Consortium Human Reference 38 (GRCh38) [33]. This leaves 2562 mutations, including 783 in the seed region. These mutations affect a total of 1209 mature miRNAs. Having filtered the databases in this manner, we focussed on mutations in the mature miRNAs outside the seed region in order to separate the effect of secondary structure from that of complementarity to the mRNA target site. We furthermore only consider point substitutions and not insertions or deletions in order to focus on mutants that have the same length as their respective WT species; this leaves 1693 mutations in 938 miRNAs. 1105 of these are associated with cancer and concern 675 miRNAs, whereas 588 are associated with other diseases and affect 400 miRNAs. Some of these entries refer to the same mutation, or different mutations that result in the same miRNA sequence, which is why the total number of unique mutated sequences is 1526 for miRNASNP-v3. The unique cancer-related mutated sequences are 1013, while the ones related to other traits and diseases are 564 in number.

In some of the analysis that follows, we consider filtered subsets of the data with different minimum numbers of mutations in the seed region or the non-seed region, as the presence of several disease-associated mutations in a given miRNA suggests a higher likelihood that the miRNA is directly involved in the disease mechanism.

Since miRNA regulate gene expression by binding to mRNAs, one may expect that mutations which alter the probability that a particular miRNA is unfolded may also affect its propensity to bind, as an unfolded molecule may interact more readily with its target than a stably folded one. In particular, one may expect mutations that lower the likelihood that a miRNA is unfolded to affect miRNA function and thus be associated with disease. We calculated the changes in the probability of the unfolded state with respect to the WTs in miRNASNP-v3 and compared them with the other possible mutations outside the seed region. We use four metrics to measure changes in RNA secondary structure as a result of mutations. We outline these metrics below.

### Change in the probability of the fully unfolded state

We calculate the Boltzmann probabilities *p*_unfolded,WT_ and *p*_unfolded,mutant_ that WT and mutant miRNAs are fully unfolded, using the ViennaRNA program RNA-subopt (see Methods). We then consider the difference between these two quantities

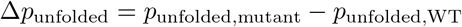

for all mutants in the point-mutational neighborhood of the WT except those that have an altered seed region and rank them by Δ*p*_unfolded_. We establish whether Δ*p*_unfolded_ is a useful predictor of the association between mutations and disease by ranking all mutants in *descending order* of their values of Δ*p*_unfolded_ and building a receiver operating characteristic (ROC) curve [34], which indicates whether disease-related mutants tend to rank higher than the rest (Figure 2).

**Figure 2:**
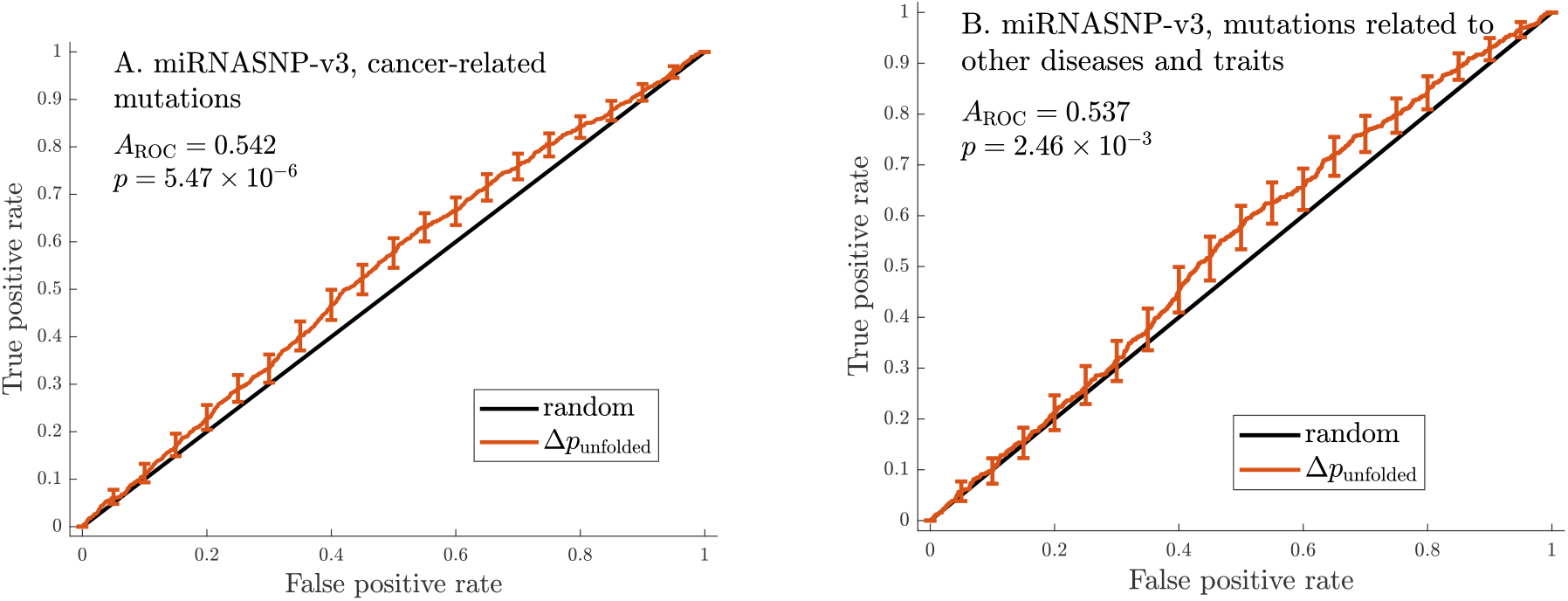
The change in probability that the miRNA is fully unfolded associated with a mutation is a predictor of the mutation’s relationship with disease. ROC curves built using Δ*p*_unfolded_ = *p*_unfolded mutant_ −*p*_unfolded WT_ as the criterion for predicting disease-related mutations. Curves based on all data on cancer-related mutations (A) and mutations related to other traits and diseases (B) from miRNASNP-v3, either with no filtering based on *N*_mut seed_ and *N*_mut non−seed_. Error bars indicate pointwise standard deviations calculated at 21 equally spaced points with the bootstrapping method [36]. The area under the ROC curve (*A*_ROC_) and the Mann-Whitney *p*-value [37] for the curves are also indicated. The Δ*p*_unfolded_ criterion performs significantly better than the random one for all both datasets (*p <* 0.05), indicating that the probability that disease-related mutants are fully unfolded tends to be higher than that for other mutants. This could be because mutants with a higher *p*_unfolded mutant_ have a higher activity than their respective WTs, and, in the case of cancer-associated mutations, they may be more effective at downregulating tumour suppressor genes.

### Change in the secondary structure Boltzmann ensemble

If the folding of a miRNA plays a role in its function, then disease-related mutations can be expected to be the ones that cause the greatest change in miRNA secondary structure. We formulate quantitative measures of the phenotypic distance of mutants from their WTs, rank the mutants in *descending order* according to these measures, i.e., compile a ranking with mutants that are furthest away from the WT at its top. Since our hypothesis is that miRNA SS impacts function, we expect mutants that rank high according to these criteria to have a stronger association with disease than other mutants. We quantify the difference between two secondary structure Boltzmann ensembles by calculating the average Hamming distance across all pairwise comparisons between structures in the two different ensembles (see Methods). We then use the average mutant-WT Hamming distance ⟨*d*_Hamming_⟩ to formulate a coarse-grained criterion, namely the percentile ranking of each individual mutant within its point-mutational neighborhood. We build ROC curves for this criterion based on data from miRNASNP-v3, whether the corresponding Mann-Whitney *p*-values fall below our significance threshold of 0.05.

### Change of the positional entropy of the mutated site

For any given site in an RNA molecule, we can calculate the positional entropy *S*^(i)^, which measures how variable the pairing of this site is within the Boltzmann ensemble. A site with low *S*^(i)^ is thus one that is consistently paired or unpaired across the ensemble, whereas a high *S*^(i)^ indicates that the site participates in pairings in many structures, but is unpaired in many others. We quantify this effect for all mutants in the partial point mutational neighborhoods of the miRNAs represented in miRNASNP-v3 by calculating the difference between the positional entropy of the mutated site for mutant and the WT

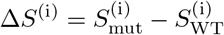

as a measure of this change in stability (see Methods).

### Change of the average positional entropy

If the secondary structure of a microRNA affects its interaction with its targets, then one may hypothesize that the mutations which significantly change the stability of that structure would be the ones with the greatest effect on miRNA function. We use the difference between the average positional entropy for the mutant and the WT, ⟨Δ*S*⟩ = ⟨*S*_mut_⟩−⟨*S*_WT_⟩ as a measure of this change in stability (see Methods). We calculate ⟨*S*_mut_⟩ for the point mutational neighborhoods of the WT miRNAs represented in miRNASNP-v3, while keeping the sequences of their seed regions (sequence positions 2-7) fixed. The positional entropy of a fold is a measure of its stability, with low ⟨*S*_mut_⟩ indicating a stable secondary structure [35]. As a miRNA needs to bind to an mRNA in order to regulate gene expression, one may expect that the stability of its fold is relevant to its function. In particular, one may expect that mutants that are more stably folded (i.e., have lower ⟨*S*_mut_⟩) than the respective WT may be less effective at binding to mRNA and thus gene regulation, potentially leading to disease.

The secondary structure of an miRNA may have an effect on its function in two ways - it could either have its own functional purpose, or it could affect the interaction with the target site as it would make the binding sites within the miRNA less accessible. In the first case, a mutant may disrupt function by causing a change to the MFE secondary structure or making it less stable; in the second one, it a mutation may interfere with miRNA function by causing sites essential to target binding to enter base pairs. The four metrics outlined above aim to quantify different types of secondary structure changes in order to detect any kind of association between modified miRNA secondary structure and disease.

## 2 Results

### 2.1 A significant proportion of disease-related miRNA mutations are associated with secondary structure changes

In Table 1, we compare the quantitative measures outlined above in terms of their power to predict disease-related mutations. To illustrate the performance of the criteria, we present the underlying ROC curves for Δ*p*_unfolded_ in Figure 2. We apply the three criteria to data from miRNASNP-v3, split between mutations associated with cancer and other traits and diseases. We provide additional ROC curves in the Supplementary Information, where we give details of other criteria we considered.

**Table 1:**
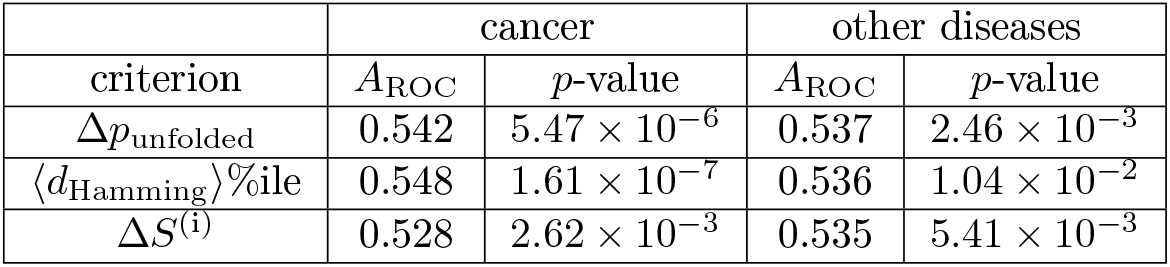
The three main criteria for measuring changes in miRNA secondary structure as a result of mutations are Δ*p*_unfolded_, the ⟨*d*_Hamming_⟩ percentile, and Δ*S*^(i)^. Mutations are ranked by these criteria, and the receiver-operator characteristic *A*_ROC_ is used to establish whether they predict disease-associated mutations in the miRNASNP-v3 dataset, split into mutations associated with cancer and those with other diseases. The *A*_ROC_ values exceed 0.5 significantly for all three metrics in both subsets of the data, suggesting that at least for some miRNAs secondary structure changes can be indicative of disease association. When checking for significance, we calculate the Mann-Whitney *p*-value and set the threshold to 0.05

|  | cancer |  | other diseases |  |
| --- | --- | --- | --- | --- |
| criterion | $A_{\text{ROC}}$ | $p$-value | $A_{\text{ROC}}$ | $p$-value |
| $\Delta p_{\text{unfolded}}$ | 0.542 | $5.47 \times 10^{-6}$ | 0.537 | $2.46 \times 10^{-3}$ |
| $\langle d_{\text{Hamming}} \rangle \%$ile | 0.548 | $1.61 \times 10^{-7}$ | 0.536 | $1.04 \times 10^{-2}$ |
| $\Delta S^{(i)}$ | 0.528 | $2.62 \times 10^{-3}$ | 0.535 | $5.41 \times 10^{-3}$ |

Δ*p*_unfolded_ is significantly better at predicting disease-associated mutations than the random criterion for data from miRNASNP-v3, suggesting that miRNAs with a fully unfolded mature form differ in activity from folded ones. In particular, disease-related mutations tend to increase the likelihood that a miRNA is unfolded more than other mutants. We show the ROC curves for this criterion for the two subsets of mutations from miRNASNP-v3 in Figure 2.

Using the percentile ranking of ⟨*d*_Hamming_⟩ of mutants within their neighborhoods also leads to *A*_ROC_ significantly greater than 0.5 for data from miRNASNP-v3. This is an independent indication that miRNA secondary structure plays a role in miRNA association with disease; as mutants are sorted in *ascending order* of this criterion, the analysis indicates that disease-related mutations tend to change secondary structure more than other mutations.

The criterion based on Δ*S*^(i)^, the difference in the positional entropy of the mutated site, performs better than random for the data in miRNASNP-v3. As we rank mutants in *ascending order*, this means that disease-associated mutants tend to be those for which the difference 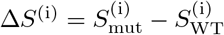 is lower.

We considered alternative approaches to quantifying secondary structure changes as well as ways to filter the data according to the number of mutations; the interested reader may find them in the Supplementary Information.

### 2.2 Disease-related mutations change the secondary structures of a specific set of individual miRNAs

We next apply the criteria formulated above to data for mutations in individual miRNAs. We use data from the miRNASNP-v3 database for both cancers and other traits and diseases, and focus on the two criteria that work best for the larger datasets - one based on the change in the probability that the miRNA is completely unfolded (Δ*p*_unfolded_) and the other based on the normalized average Hamming distance from the WT (⟨*d*_Hamming_⟩*L*^−1^). Note that at the level of the individual miRNA ⟨*d*_Hamming_⟩*L*^−1^ and the miRNA-specific percentile of ⟨*d*_Hamming_⟩ produce identical rankings. We observe a signal in Δ*p*_unfolded_ (*p <* 0.05) for 44 miRNAs, listed in Table 12 (see Supplementary Information) and ⟨*d*_Hamming_⟩ *L*^−1^ - for 48, listed in Table 13 (see Supplementary Information), and for thirteen miRNAs *p <* 0.05 for both criteria.

Since we observe that multiple criteria perform better than random for some miRNAs, we also combine the *p*-values obtained from independent tests using Fisher’s method [38]. We do that for ⟨Δ*p*_unfolded_⟩ and ⟨*d*_Hamming_⟩ *L*^−1^, and we give the results for miRNASNP-v3 in Table 14. We observe a signal in 46 miRNAs from this set, including seven miRNAs for which the combined *p*-value is below 0.05 while the individual Mann-Whitney *p*-values are not. Intriguingly, we observe that the distributions of *A*_ROC_ within the sets of miRNAs for which these two criteria perform better than random exhibit signs of bimodality, with clear peaks around 0 and 1 and few, if any, points in between them. We quantify this observation by calculating the bimodality coefficients of the distributions according to [39], and obtain the values 0.714 for Δ*p*_unfolded_ and 0.745 for *d*_Hamming_ *L*^−1^. Coefficient values above 5/9 suggest bimodality [39, 40].

A possible explanation of this result is that the WTs of miRNAs clustered around one peak contribute to the initiation and/or progression of disease, whereas those around the other peak are essential for disease-prevention, e.g., because they act as tumour-suppressors. In that case, if miRNA folding affects function, we would expect that the mutants most strongly associated with disease in disease-suppressing miRNAs would be those that change the secondary structure the most, which would translate into *A*_ROC_ *>* 0.5 for Δ*p*_unfolded_ and *A*_ROC_ *<* 0.5 for ⟨*d*_Hamming_⟩ *L*^−1^. The comparatively small overlap between the miRNAs for which Δ*p*_unfolded_ and ⟨*d*_Hamming_ ⟩*L*^−1^ perform significantly better than random suggests that some miRNAs need to be fully unfolded to perform their functions, whereas for others the folding itself plays a functional role.

When performing a large number of statistical tests, as we do in the current section for hundreds of miRNAs, a *p*-value of 0.05 is not a sufficient indicator of statistical significance. For this reason, in addition to testing whether *Δp*_unfolded_ fulfils the Mann-Whitney test for statistical significance for individual miRNAs, we estimate the positive false discovery rate (pFDR), which is the FDR in case there is at least one positive finding. We then calculate the *q*-value, an FDR-based measure of significance equal to the minimum positive false discovery rate at which a test with *p*-value *p*_i_ is considered significant. We compute the Benjamini-Hochberg linear step-up procedure as implemented in the built-in MATLAB R2022b function mafdr [41]. We consider various levels of filtering by *N*_mut seed_ and *N*_mut non−seed_ since we expect that a large number of recorded disease-related mutations for a particular miRNA implies a stronger association between the latter miRNA and disease. Moreover, focussing on a smaller number of miRNAs makes it possible to obtain lower *q*-values as it requires fewer tests. The number of unique sequences with mutations outside the seed is 452 at *N*_mut seed_ ≥ 1 and *N*_mut non−seed_ ≥ 1, 172 at *N*_mut seed_ ≥ 2 and *N*_mut non−seed_ ≥ 2, and 113 at *N*_mut seed_ ≥ 3 and *N*_mut non−seed_ ≥3.

Using a *q*-value threshold of 0.05 we find five miRNAs (given in Table 2) for which the association between disease-related mutations and secondary structure changes is significant (in one case borderline). We employ two different filtering regimes, the first requiring at least one mutation in the seed region and non-seed region (*N*_mut seed_, *N*_mut non−seed_ ≥ 1), and the second requiring at least three mutations in each region (*N*_mut seed_, *N*_mut non−seed_ ≥3). The former covers a wider range of mutants, but this means a higher bar for significance when using the Benjamini-Hochberg procedure. The latter focuses on a smaller number of miRNAs that appear to have many disease-related mutations and are thus likely to be disease-associated. Figure 3 is an illustration of the data underlying Table 2 for individual miRNas with the tumour suppressor hsa-miR-4537 as an example. The figure contains ROC plots characterizing the performance of Δ*p*_unfolded_ and ⟨*d*_Hamming_⟩ *L*^−1^, both of which have an associated *p*-value of less than 0.05. We use the Fisher method to calculate a combined *p*-value for the two criteria and apply the Benjamini-Hochberg procedure to calculate the minimum false discovery at which the results are significant, i.e., the *q*-value. Applying a more stringent filter by the number of reported disease-related mutations in the mature region decreases the *q*-value because it restricts the set of miRNAs whose *p*-values are processed by the Benjamini-Hochberg method, excluding miRNAs with few mutations for which the uncertainty is greater.

**Table 2:** Benjamini-Hochberg *q*-values, i.e., minimum positive false discovery rates, based on combined Mann-Whitney *p*-values for Δ*p*_unfolded_ and ⟨*d*_Hamming_⟩ *L*^−1^. Significant values (*q*-value *<* 0.05) are emphasised in bold. The first two miRNAs (hsa-miR-4641 and hsa-miR-1269b) have fewer than three miRNASNP-v3 entries for mutations in the seed region or the non-seed region, and are thus excluded by the more stringent filter (*N*_mut seed_, *N*_mut non−seed_ ≥3). The last miRNA, hsa-miR-6821-3p, is borderline significant, but is nevertheless mentioned here because it becomes fully significant (*q* = 2.996 × 10^−2^) if we combine the *p*-values for Δ*p*_unfolded_ and ⟨*d*_Hamming_⟩ *L*^−1^ with that for Δ*S*^(i)^. All four miRNAs have been linked with disease [42–48]. Note that two pairs of miRNAs have equal *q*-values. The reason for this is that the *p*-values for mutants come from a small discrete set, which can yield the same *q*-value in the Benjamini-Hochberg procedure that ranks them in ascending order and multiplies them by a factor involving their rank.

| miRNA | filtering levels |  |
| --- | --- | --- |
| | $N_{\text{mut seed}}, N_{\text{mut non-seed}} \geq 1$ | $N_{\text{mut seed}}, N_{\text{mut non-seed}} \geq 3$ |
| **hsa-miR-1269b** | **$2.666 \times 10^{-2}$** | - |
| **hsa-miR-4477b** | $6.792 \times 10^{-2}$ | **$1.313 \times 10^{-2}$** |
| **hsa-miR-4537** | $6.792 \times 10^{-2}$ | **$1.864 \times 10^{-2}$** |
| **hsa-miR-4641** | **$2.666 \times 10^{-2}$** | - |
| **hsa-miR-6821-3p** | $> 0.3$ | $6.299 \times 10^{-2}$ |

**Figure 3:**
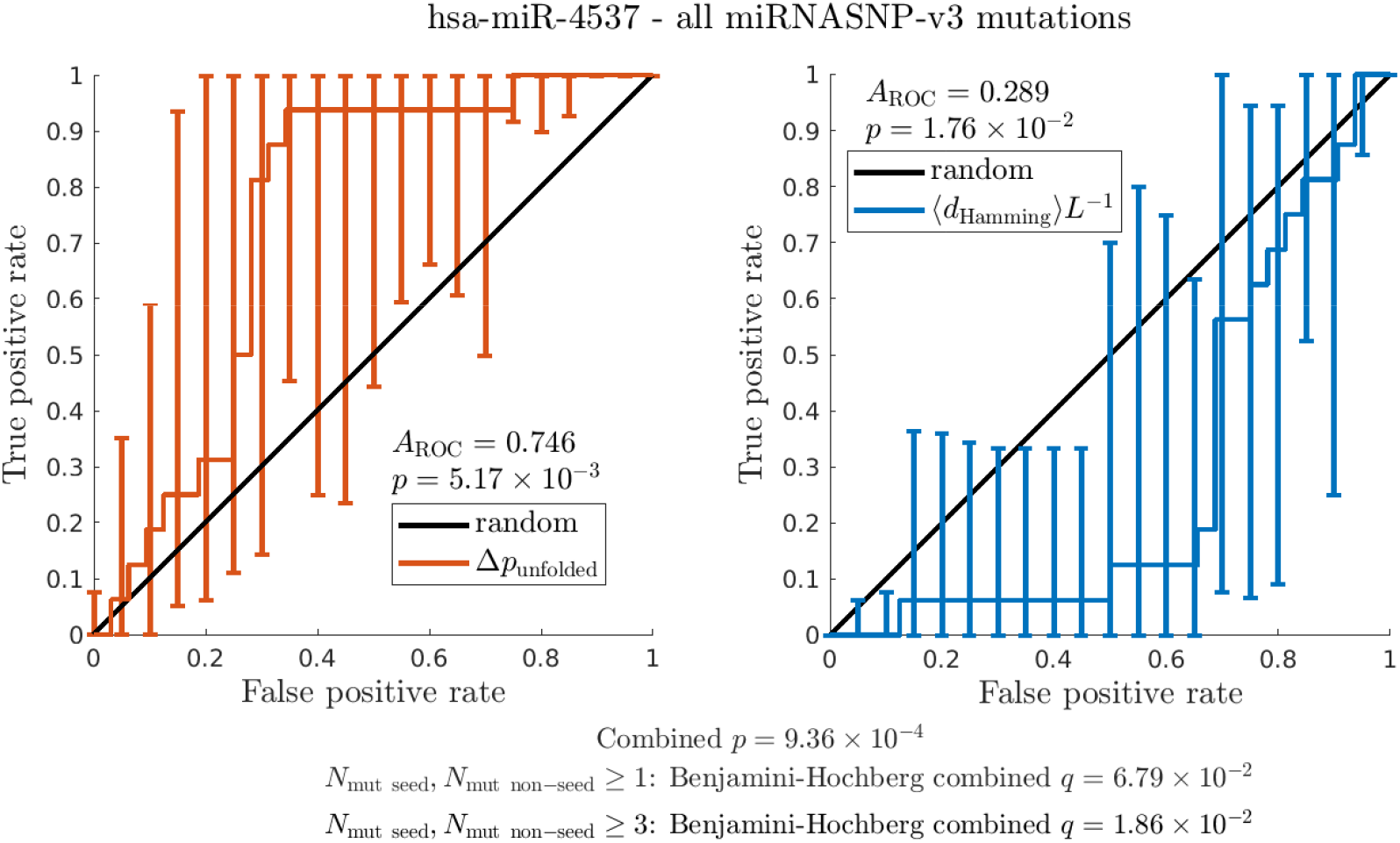
Performance of Δ*p*_unfolded_ and ⟨*d*_Hamming_⟩ *L*^−1^ as predictors of disease association for mutations in the tumour suppressor hsa-miR-4537, which is known to be relevant to gastric cancer [46]. Both of these metrics for the effect of point mutations on miRNA secondary structure perform significantly better than random when applied to the point-mutational neighborhood of hsa-miR-4537. Note also that ⟨*d*_Hamming_⟩ *L*^−1^ is equivalent to the percentile-based measure for aggregated datasets.

**Figure 4:**
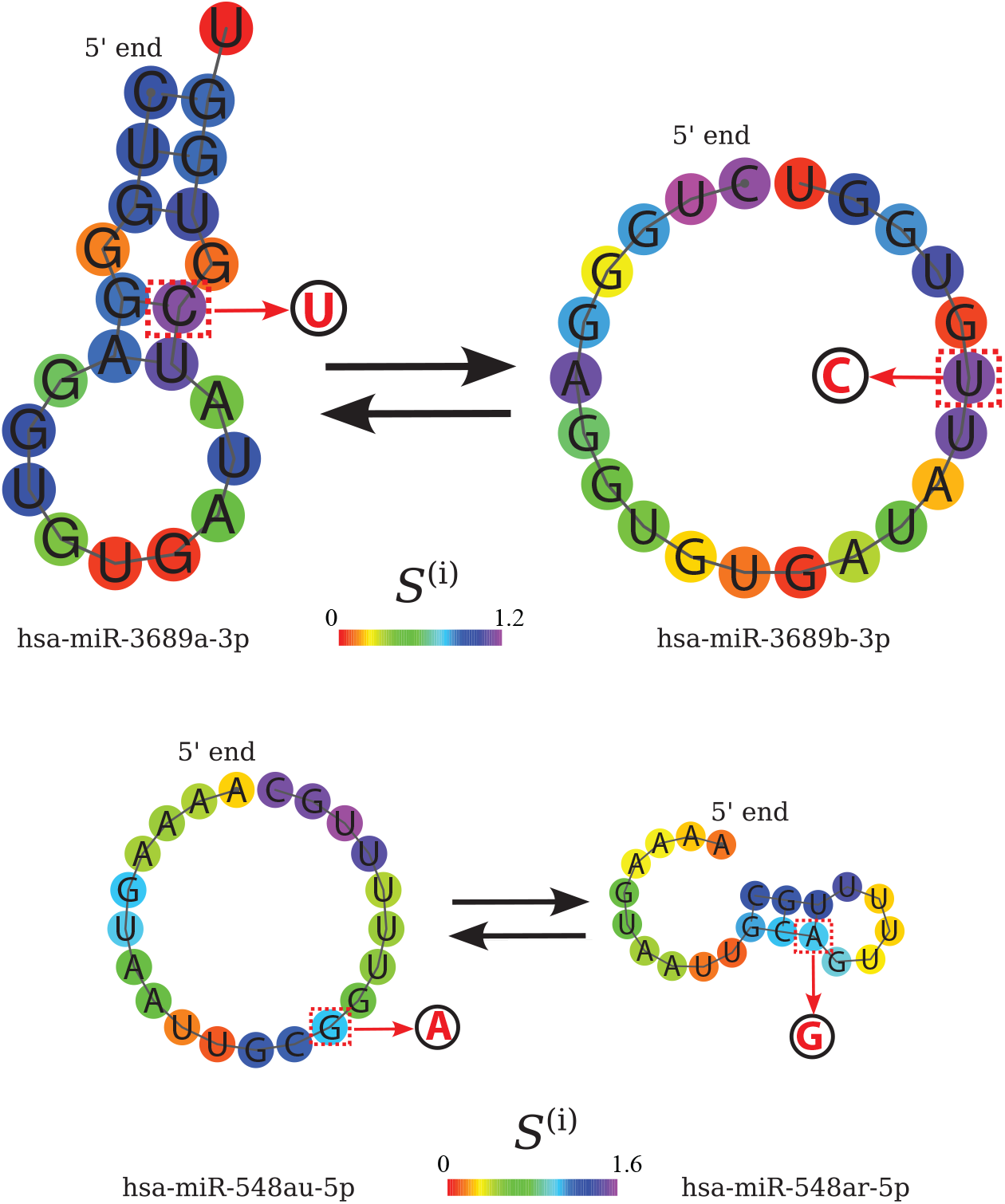
Single point substitution mutations that convert one miRNA to another. Top: substituting the C in position 17, which has the highest predicted positional entropy (see color and colorbar) in the MFE structure, changes hsa-miR-3689a-3p into hsa-miR-3689b-3p. As the mutation converts a Watson-Crick GC pair into a GU wobble pair, the MFE structure changes from stably folded to fully unfolded. Both the forward and the reverse mutations are associated with disease in miRNASNP-v3. Bottom: substitution of G in position 13 with A creates a AU Watson-Crick pair, changing the MFE structure from fully unfolded to stably folded. These MFE predictions and visualizations are based on tools from the ViennaRNA suite [50].

Of the five miRNAs, three (hsa-miR-1269b, hsa-miR-4537 and hsa-miR-4477b) are known to play a role in cancers: hsa-miR-1269b is associated with hepatocellular carcinoma [42–44], and a mutation in the gene coding for hsa-miR-4477b has been reported to occur during the transformation of colorectal adenoma into colorectal cancer [45], and hsa-mir-4537 is a tumour suppressor in gastric cancers [46]. hsa-miR-6821-3p has been reported to be an extracellular genomic biomarkers of osteoarthritis [47], and a potential biomarker for cancer [48].

The fact that only a small number of individual miRNAs are highlighted by this approach is partly due to the fact that the analysis places a lower bound on the *p*-values. This is because the relatively small number of disease-associated mutations outside the seed region (typically one or two, versus 48 in a typical neighbourhood) means that for many miRNAs, the lowest possible *p*-values are 4.17 × 10^−2^ (1 mutant) or 1.57 × 10^−3^ (2 mutants).

### 2.3 Disease-related point mutations may convert one miRNA WT into another

Interestingly, the miRNASNP-v3 database contains several point mutations that convert one miRNA WT into another. A striking example is the mutation of C at position 17 to G, which turns **hsa-miR-3689a-3p** to **3689c** or **3689b-3p**, the latter two having the same mature sequence. This mutation, which occurs at the site with the highest positional entropy, converts a GC Watson-Crick pair to a wobble pair and turns the relatively stable MFE fold of hsa-miR-3689a-3p to a fully unfolded structure. Position 17 is outside the seed region and the supplementary region between nt 13-16 that is known to contribute to target recognition for some miRNAs [10, 11]. Thus, the mutation probably does not affect the miRNA-mRNA target complementarity and it appears likely that the drastic change in secondary structure affects miRNA activity. Moreover, the reverse mutations, which change hsa-miR-3689c or 3689b-3p to hsa-miR-3689a-3p are also reported to be associated with disease in miRNASNP-v3. miRNAs from the hsa-mir-3689 family have been shown to be differentially expressed in conjunctival malignant melanoma, a rare form of cancer [51].

Another mutation (ID: rs138894217) that converts one miRNA into another and dramatically changes its fold is the substitution of A to G at position 13 in hsa-miR-548ar-5p, which turns it into hsa-miR-548au-5p. It turns a AU Watson pair into GU wobble pair, making the MFE structure fully unfolded in contrast with the stably folded WT. Although mutation rs138894217 is associated with heel bone mineral density and body mass index according to miRNASNP-v3, we were not able to find information about it in the references cited therein. This points to another possible confounder in our study - some of the associations with disease that we attempt to predict here may be simply due to mistakes in the database we source them from.

A disease-associated mutation mutation of G at site 13 in **hsa-miR-1269a** to A, which changes a UG wobble pair to a UA Watson-Crick pair, changes the stability of the MFE structure and turns the sequence

As in the case of aggregated datasets with mutations for many miRNAs, mutations are ranked by these criteria, and the area under the receiver-operator characteristic *A*_ROC_ is used to establish whether disease-associated mutations cluster at one end of the ranking. The pointwise confidence bounds for the true positive rates in the ROC plots and the confidence bounds for *A*_ROC_ are calculated via bootstrapping and vertical averaging for 21 equally spaced values of the false positive rate from 0 to 1. The confidence interval is computed through bootstrapping via the bias corrected and accelerated percentile method, as implemented in the MATLAB function bootci; for more details, see [49] and the references cited therein. The *A*_ROC_ are significantly different from the value for the random criterion (0.5) for both of these metrics. This suggests that secondary structure changes can be indicative of disease association for this particular miRNA, and, in agreement with this conclusion, combining these Mann-Whitney *p*-values via the Fisher method yields *q*-values (minimum positive false discovery rates) of less than 0.05 when subjected to multiple hypothesis testing, see Table 2.

According to both criteria, mutations that introduce a greater change in secondary structure tend to be associated with disease. This results in *A*_ROC_ *<* 0.5 for ⟨*d*_Hamming_⟩ *L*^−1^ because, for consistency with Table 1, we rank mutations in *ascending* order of their associated ⟨*d*_Hamming_⟩ *L*^−1^, see the Methods section. This means that disease-associated mutations whose secondary structure tend to have a greater SS Hamming distance from the WT than other mutations for this individual miRNA, but a smaller one for the aggregated datasets. into that of **1269b**. While the mutant and the WT have the same fold, the ensemble is substantially changed, with a normalized average Hamming distance between the mutant and the WT of ⟨*d*_Hamming_⟩ *L*^−1^ = 0.35. Site 13 is in the supplementary region and may contribute to mRNA target recognition. This particular mutation (rs73239138), is known to be associated with various types of cancer, e.g., hepatocellular carcinoma [42–44].

We provide data on all mutations of this type that we have identified in Table 15, which contains information on the WT and mutated sequences and SS, as well as the ⟨*d*_Hamming_⟩ *L*^−1^ and Δ*p*_unfolded_ for them.

## 3 Discussion and conclusions

This computational study explores the role of secondary structure in mature miRNA mutants based on literature data for associations between such mutants and disease. We formulate different quantitative measures for the difference between a given mutant and its respective WT and test their ability to predict disease association.

Our results for the data in miRNASNP-v3 indicate a significant association between disease and mutations that increase the probability that a given mature miRNA is unfolded. This is in contrast to the work of Diederichs and Haber [18], who found that mutations which significantly changed pri-miRNA secondary structure had no effect on their processing and maturation. Moreover, we see a significant effect of the change in fold stability caused by mutations across the miRNASNP-v3 database. When we apply our criteria for mutation-induced secondary structure changes to the data for individual miRNAs, we observe a particularly strong relationship between secondary structure and disease for several miRNAs. This analysis combines *p*-values for the (*Δp*_folded_) and (⟨*d*_Hamming_⟩ */L*) metrics using the Fisher method and employs the Benjamini-Hochberg method for multiple hypothesis testing. We furthermore filter the data according to the number of mutations recorded in the seed region and outside the seed region. The miRNAs that emerge are **hsa-miR-1269b, hsa-miR-4537, hsa-miR-4477b, hsa-miR-4641**, and **hsa-miR-6821-3p**, which include three miRNAs that are known to play a role in cancers [42–46]. It is likely that the significance criteria are not met for many other miRNAs because of a combination of factors: 1) the small number of mutations recorded for most miRNAs and 2) the different nature of the mutations in the database. Most of the disease-associated mutations in miRNASNP-v3 have been identified via methods such as genome-wide association studies (GWAS), which do not establish a causative relationship with disease, meaning that some of some these mutations are likely to simply accompany disease. Another possible confounder in our current study are mistakes in the assignment of disease association to mutations, of which we give an example in the previous section (SNP rs138894217). Quantitative experimental data on the activity of various point mutants of a set of miRNAs would shed light on these matters.

As we note in the introduction, it is difficult to establish the role of mature miRNA SS because the mature form of the molecule is incorporated in RISC and, furthermore, even mutations in the mature sequence could affect miRNA function through changing the folding and maturation of its precursor, rather than through changing the SS of the mature miRNA itself, see the example of miRNA-125a in Ref. [22] and others in Ref. [52]. With all this in mind, a definitive test of our hypothesis that miRNA SS can have significant influence on function requires experimental data on the regulatory activity of mutants, preferably in the form of gene suppression activity for the point mutants derived from several mature miRNAs (with the seed region fixed). The availability of such experimental data would allow for the further refinement of our methods for predicting which mutations are associated with disease, particularly cancer, and thus yield a potentially valuable diagnostic tool.

## 4 Methods

### Secondary structure prediction

We used version 2.5.0 of the ViennaRNA package [50] to predict and analyze the secondary structure of mutant and WT miRNAs from the two filtered datasets.

### Statistical significance of ROC results

We use the area under the ROC curves, *A*_ROC_, as a measure of the power of Δ*p*_unfolded_ as a predictor of the association of a given mutation with disease. We then assess the statistical significance of the results by calculating the 95% confidence intervals for *A*_ROC_ via the bootstrapping method as implemented in MATLAB R2022b [36]. Furthermore, for cases in which the 95% confidence interval excludes the *A*_ROC_ for a random predictor, 0.5, we calculate the two-sided Mann-Whitney *p*-values for the null hypothesis that the Δ*p*_unfolded_ distributions for the disease-related and other mutants have the same median; we use the built-in MATLAB R2022b function for this purpose as well [37]. The pointwise confidence bounds for the true positive rates in the ROC plots and the confidence bounds for *A*_ROC_ are calculated via bootstrapping and vertical averaging for 21 equally spaced values of the false positive rate from 0 to 1. The confidence interval is computed through bootstrapping via the bias corrected and accelerated percentile method, as implemented in the MATLAB function bootci; for more details, see [49] and the references cited therein.

### Boltzmann frequency of structures

The frequency of a particular secondary structure *s* with energy *E*(*s*) in the thermodynamic equilibrium ensemble is [53]

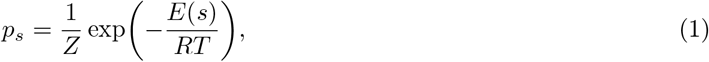

with *R* denoting the gas constant, *T* - the temperature, for which we use the biologically relevant value 310.15 K, and 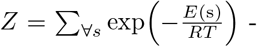 the partition function; for the fully unfolded state, there is no base-pairing, *E*(*s*) = 0, and *p*_unfolded mut_ = *Z*^−1^.

### RNAsubopt

We use RNAsubopt to generate a sample of structures drawn randomly from the Boltzmann ensemble according to their probability, starting with a sample size of 10. In case the fully unfolded structure is not present in the initial sample, we increase the sample size by a factor of 10 and we repeat the process until we either encounter the trivial SS or reach 10^10^ samples. In the latter case, the Boltzmann weight for this structure is negligible and we use the approximation *p*_unfolded_ ≈0.

### Hamming distance metric

We measure the differences between two secondary structures *s*_1_ and *s*_2_ in the dot-bracket representation, where dots represent unpaired sites and brackets - base pairs see e.g. [54], via the Hamming distance between the two, i.e., the number of sites for which there are discrepancies (*d*_Hamming_(*s*_1_, *s*_2_)). For each mutant in the partial point mutational neighborhoods of the miRNAs in miRNASNP-v3, we average the pairwise Hamming distance between the possible secondary structures in its equilibrium ensemble and those for the equilibrium ensemble of the WT, weighted by their joint Boltzmann probability,

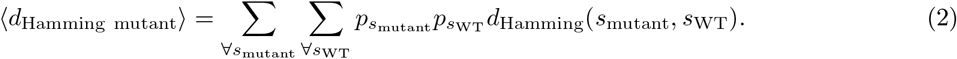

The sums in Eq. (2) contain the secondary structures that RNAsubopt has drawn from the respective equilibrium ensembles on the basis of a sample with 10^5^ entries. Structures that have not been encountered in drawing this sample have only a negligible contribution to ⟨*d*_Hamming_⟩. For the criteria based on ⟨*d*_Hamming_⟩ *L*^−1^, unlike most other quantities we study, we rank mutants in *ascending order*, which means that top-ranked entries are closest to the WT. We do this to ease the readability of the results because for large datasets, mutants associated with disease tend to be closer in SS to the WT than other mutants; however, this trend is reversed for some individual point mutational neighborhoods such as that of the microRNA hsa-miR-4537, which is illustrated in Figure 3.

### Positional entropy

The positional entropy of site ‘i’ within an RNA molecule is defined as

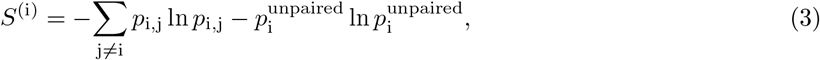

where *p*_i,j_ is the probability that site ‘i’ is paired with site ‘j’, and 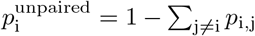 is the probability that site ‘i’ is unpaired [55]. When building ROC curves for *S*^(i)^, we sort mutants in *ascending order*, as we do for the criteria based on the mutant-WT Hamming distance. This means that the top-ranked entries are those for which the difference in positional entropy at the mutated site is lowest.

Calculating the average positional entropy, ⟨Δ*S*⟩, for an RNA molecule from the positional entropies of the individual sites, *S*^(i)^, requires straighforward application of the definition in Eq. (3)

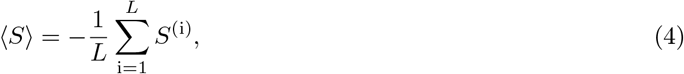

where *L* is the number of nucleotides in the RNA molecule. The values of ⟨*S*⟩ for the mutant and the WT are comparable only if 1) the minimum-free-energy (MFE) structures both of them have secondary structure or 2) neither MFE structure has secondary structure. Note that this latter case is much less common as base-pairing is energetically favourable and usually possible. We analyze these two cases separately; as the criterion based on the ⟨Δ*S*⟩ for non-folded mutants and WTs performs no better than the random criterion, we only discuss it in the Supplementary Information, which also contains more detail on the definition of the average positional entropy.

## 5 Acknowledgements

This work was supported by the Isaac Newton Trust (NQAG/341) and was performed using resources provided by the Cambridge Service for Data Driven Discovery (CSD3) operated by the University of Cambridge Research Computing Service (www.csd3.cam.ac.uk), provided by Dell EMC and Intel using Tier-2 funding from the Engineering and Physical Sciences Research Council (capital grant EP/T022159/1), and DiRAC funding from the Science and Technology Facilities Council (www.dirac.ac.uk). JKN acknowledges funding from the MRC (Grant MC_FE_00035) Cross Disciplinary Fellowship (XDF) Programme.

