## Supplementary Information for "Changes in miRNA secondary structure can predict mutations associated with cancer and other diseases"

**3 Tables and graphical comparisons of measures of criteria performance** **12**

**4 Tables of $p$-values and other data for individual miRNAs** **12**

**5 Table of disease-associated mutations that convert one WT miRNA to another** **12**

**6 Case studies** **36**
6.1 hsa-miR-4537 . . . . . . . . . . . . . . . . . . . . . . . . . . . . . . . . . . . . . . . . . . . . 36
6.2 hsa-miR-485-5p . . . . . . . . . . . . . . . . . . . . . . . . . . . . . . . . . . . . . . . . . . . 36

**7 Additional information on miRNAs with $q < 0.05$** **41**

8 Code and data 41

Here we provide information on the criteria for relationship with disease that we only mentioned in the main text, additional ROC curves for the different predictors, plots that compare $A_{\mathrm{ROC}}$ for different criteria under different levels of filtering, and tables with $A_{\mathrm{ROC}}$ and $p$-values and sample sizes.

### 1 SomamiR database

In addition to the miRNASNP-v3 dataset discussed in the main text, we evaluate the performance of our metrics on the SomamiR dataset of miRNA mutations encountered in cancer [1]. After cross-checking the SomamiR data with HG38 and miRBase, the number of mature miRNA mutations outside the seed recorded in it is 439, with 363 unique sequences derived from 285 WT miRNAs, and the number of seed-region mutations is 173. Filtering to include only miRNAs with $N_{\mathrm{mut\ seed}} \geq 1$ and $N_{\mathrm{mut\ non-seed}} \geq 1$ leaves 81 mutations outside the seed region, representing 66 unique sequences and affecting 40 miRNAs.

### 2 Additional metrics of the effect of mutations on secondary structure

We formulated several other criteria that measure the effect of non-seed point mutations on miRNA secondary structure, and give a list of them below. We do not describe them or give details on their performance in the main text since the latter is inferior to that of the three we discuss therein.

a) The change in the probability that the seed region is fully unfolded, $\Delta p_{\mathrm{unfolded\ seed}} = p_{\mathrm{unfolded\ seed\ mutant}} - p_{\mathrm{unfolded\ seed\ WT}}$. As the seed is key for target recognition, one may expect that a high value of $\Delta p_{\mathrm{unfolded\ seed}}$ would strongly affect the activity of a miRNA.

b) The average positional entropy of a mutant, $\langle S_{\mathrm{mut}} \rangle$, which measures the stability of a fold. We defined additional criteria related to $\langle S_{\mathrm{mut}} \rangle$ as follows.

- The absolute mutant positional entropy, $\langle S_{\mathrm{mut}} \rangle$.
- The difference in positional entropy between mutant and WT, $\langle \Delta S \rangle = \langle S_{\mathrm{mut}} \rangle - \langle S_{\mathrm{WT}} \rangle$. Apart from applying this criterion to the case in which both the WT and the mutant are folded as discussed in Section 2.3, we also used it for **i)** data for all miRNAs in a dataset and **ii)** datasets filtered to only include fully unfolded WT and mutant miRNAs.

In building the ROC curves and analyzing the performance of these criteria, we use the same approach described above for $\Delta p_{\mathrm{unfolded}}$ - we calculate the criterion values for a set of mutants, then rank the latter in *descending order* according to that criterion. Next, we ascertain whether the criterion predicts association with disease better than random by building a ROC curve, calculating the area under it ($A_{\mathrm{ROC}}$) and checking whether the associated $p$-values meet our significance criteria. We provide more details on the criteria and their performance below.

#### 2.1 Change in the probability of the unfolded states with respect to the WT

##### 2.1.1 Change in the probability of the fully unfolded state with respect to the WT

Additional ROC curves for $\Delta p_{\mathrm{unfolded}} = p_{\mathrm{unfolded\ mutant}} - p_{\mathrm{unfolded\ WT}}$ are shown in Figures 1-2.

##### 2.1.2 Change in the probability that the seed region is unfolded with respect to the WT

As the miRNA seed region is crucial to miRNA-mRNA binding, we expect that mutations which change its folding have a significance in disease. Specifically, we test whether disease-related mutations change the

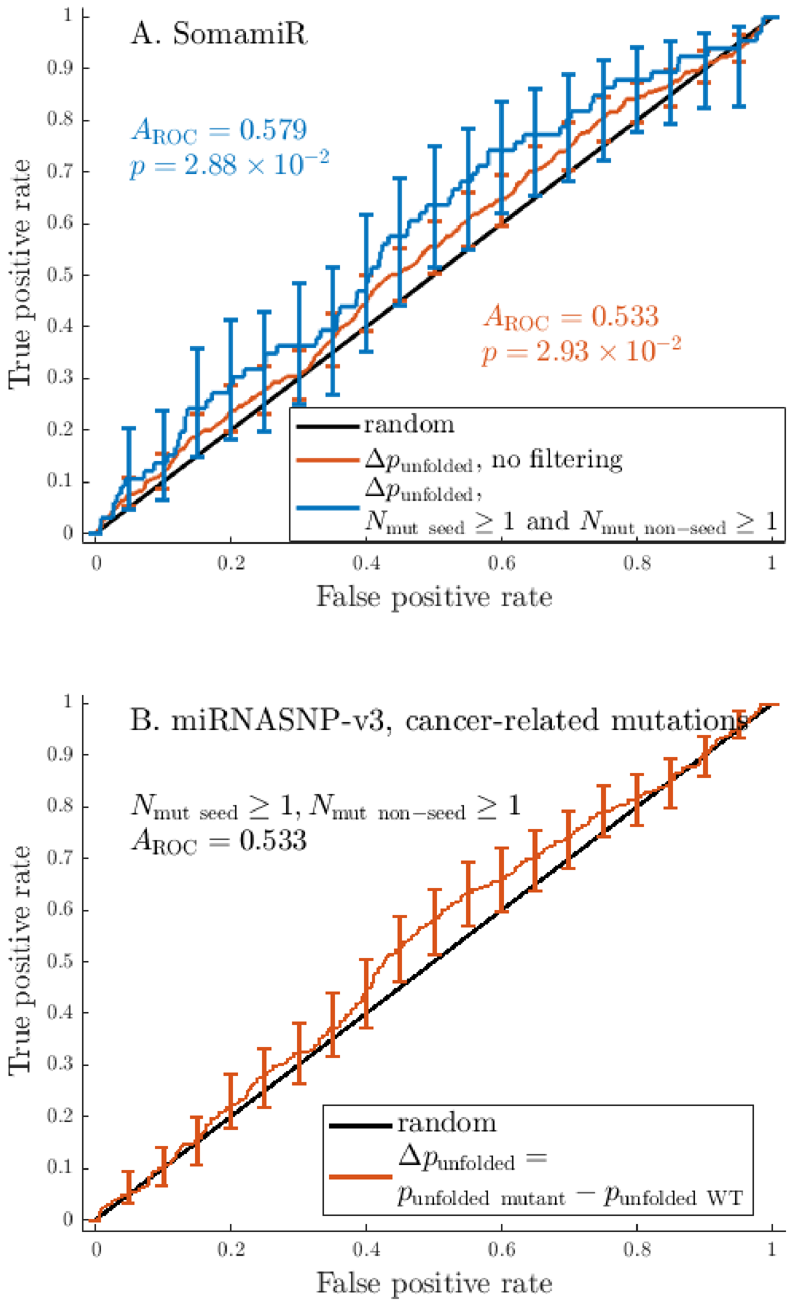

A. SomamiR
$A_{\mathrm{ROC}} = 0.579$
$p = 2.88 \times 10^{-2}$
$A_{\mathrm{ROC}} = 0.533$
$p = 2.93 \times 10^{-2}$
random
$\Delta p_{\mathrm{unfolded}}$, no filtering
$\Delta p_{\mathrm{unfolded}}$,
$N_{\mathrm{mut\ seed}} \geq 1$ and $N_{\mathrm{mut\ non-seed}} \geq 1$
True positive rate
False positive rate
B. miRNASNP-v3, cancer-related mutations
$N_{\mathrm{mut\ seed}} \geq 1$, $N_{\mathrm{mut\ non-seed}} \geq 1$
$A_{\mathrm{ROC}} = 0.533$
random
$\Delta p_{\mathrm{unfolded}} =$
$p_{\mathrm{unfolded\ mutant}} - p_{\mathrm{unfolded\ WT}}$
True positive rate
False positive rate

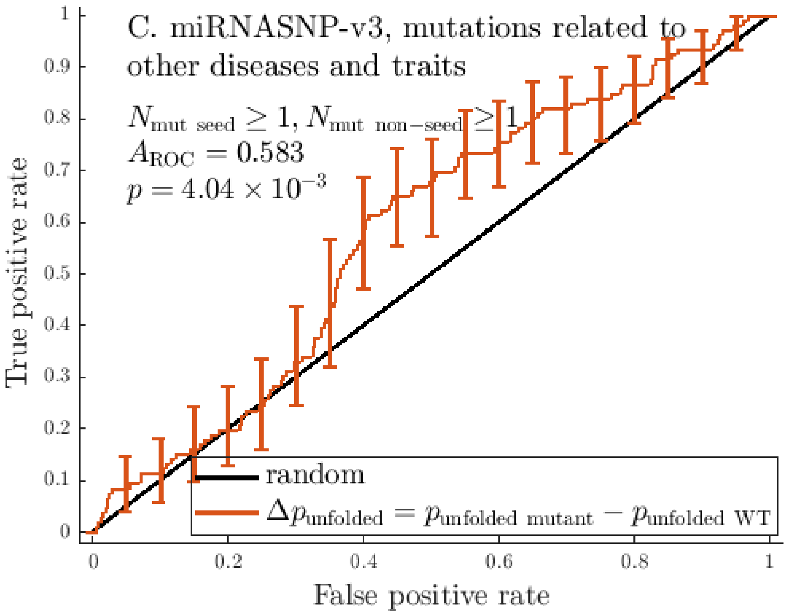

Figure 1: **The change in probability that the miRNA is fully unfolded associated with a mutation is a predictor of the mutation's relationship with disease.** ROC curves built using $\Delta p_{\text{unfolded}} = p_{\text{unfolded mutant}} - p_{\text{unfolded WT}}$ as the criterion for predicting disease-related mutations. Data from SomamiR (A), information on cancer-related mutations (B) and mutations related to other traits and diseases (C) from miRNASNP-v3, either with no filtering based on $N_{\text{mut seed}}$ and $N_{\text{mut non-seed}}$ (orange curves) or with $N_{\text{mut seed}} \geq 1$ and $N_{\text{mut non-seed}} \geq 1$ (blue curves) for all mature miRNAs considered here. Error bars indicate pointwise standard deviations calculated at 21 equally spaced points with the bootstrapping method [2]. The area under the ROC curve ($A_{\text{ROC}}$) and the Mann-Whitney $p$-value [3] for the curves are also indicated. The $\Delta p_{\text{unfolded}}$ criterion performs significantly better than the random one for all three datasets ($p < 0.05$) if no filtering is applied, indicating that the probability that disease-related mutants are fully unfolded tends to be higher than that for other mutants. This could be because mutants with a higher $p_{\text{unfolded mutant}}$ have a higher activity than their respective WTs, and, in the case of cancer-associated mutations, they may be more effective at downregulating tumour suppressor genes. When the analysis for each dataset is restricted to just the miRNAs for which at least one mutation in the seed region and the rest of the mature miRNA, $A_{\text{ROC}}$ increases and so does the certainty that the criterion outperforms the random one, except for the set of cancer-associated mutations from miRNASNP-v3, for which we observe changes in the opposite direction.

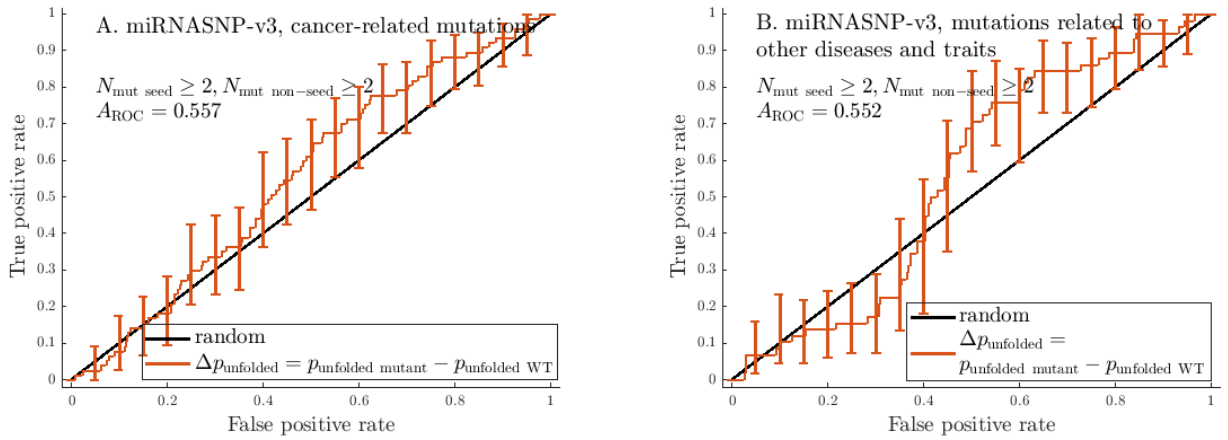

Figure 2: ROC curves built using $\Delta p_{\text{unfolded}} = p_{\text{unfolded mutant}} - p_{\text{unfolded WT}}$ as the criterion for predicting disease-related mutations. Data taken from miRNASNP-v3. Error bars indicate pointwise standard deviations calculated at 21 equally spaced points with the bootstrapping method [2]. The area under the ROC curve ($A_{\text{ROC}}$) is also indicated.

probability that the seed region contains no base-pairing more than other possible point mutations. To calculate the change in probability for no seed-region base-pairing, we impose the constraint that no bases within the seed region are paired, calculate the free energy of the constrained ensemble $F_{\text{constrained}}$ and use the equation

$$p_{\text{unfolded seed}} = \exp\left(-\frac{F_{\text{constrained}} - F_{\text{unconstrained}}}{RT}\right), \tag{1}$$

where $F_{\text{unconstrained}}$ is the free energy of the ensemble without any imposed constraints.

Having calculated $p_{\text{unfolded seed}}$ for all mutants in the point-mutational neighborhood of the WT except those that have an altered seed region, we then calculate $\Delta p_{\text{unfolded seed}} = p_{\text{unfolded seed mutant}} - p_{\text{unfolded seed WT}}$. We then use this quantity as a criterion for ranking these mutants and build an ROC curve in order to assess its power to predict disease-related mutations (Figure 3).

The change in the probability that the seed region is unfolded, $\Delta p_{\text{unfolded seed}}$, does not perform significantly better than random for any of the datasets that we studied, suggesting that the likelihood that the seed region is unfolded does not play an important role in determining miRNA activity.

#### 2.2 Hamming-distance-based criteria

##### 2.2.1 Normalized Hamming distance from the WT, $\langle d_{\text{Hamming}} \rangle L^{-1}$

We formulate another criterion based on the average Hamming distance $\langle d_{\text{Hamming mutant}} \rangle$ from the WT which registers smaller differences in secondary structure than the percentile-based one discussed in the main text. To do this, we normalize the average Hamming distance by the miRNA length $L$, $\langle d_{\text{Hamming}} \rangle L^{-1}$. We then test whether disease-related mutants from SomamiR and miRNASNP-v3 tend to change miRNA secondary structure more than other mutants by ranking mutants in *ascending order* of their $\langle d_{\text{Hamming mutant}} \rangle$ and building an ROC curve for this criterion (Figure 4).

The normalized Hamming distance, $\langle d_{\text{Hamming}} \rangle L^{-1}$, has $A_{\text{ROC}}$ significantly smaller than 0.5 for the 45 miRNASNP-v3 entries related to diseases other than cancer for which $N_{\text{mut seed}} \geq 1$ and $N_{\text{mut non-seed}} \geq 1$. Since we order mutants in ascending order of $\langle d_{\text{Hamming}} \rangle L^{-1}$, this means that mutants with a larger $\langle d_{\text{Hamming}} \rangle L^{-1}$, which rank lower, tend to be associated with disease.

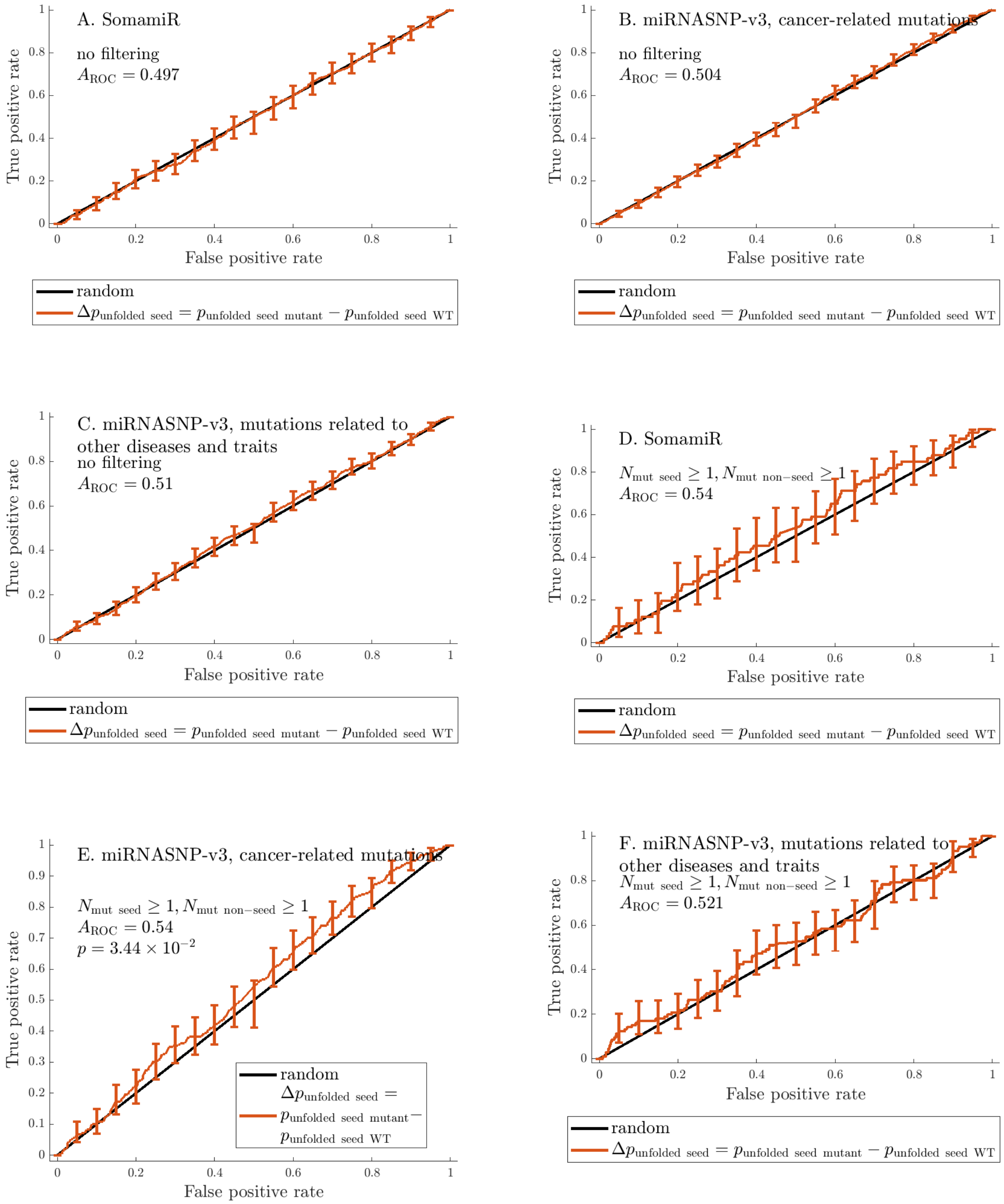
A. SomamiR
no filtering
$A_{\text{ROC}} = 0.497$
B. miRNASNP-v3, cancer-related mutations
no filtering
$A_{\text{ROC}} = 0.504$
C. miRNASNP-v3, mutations related to other diseases and traits
no filtering
$A_{\text{ROC}} = 0.51$
D. SomamiR
$N_{\text{mut seed}} \geq 1, N_{\text{mut non-seed}} \geq 1$
$A_{\text{ROC}} = 0.54$
E. miRNASNP-v3, cancer-related mutations
$N_{\text{mut seed}} \geq 1, N_{\text{mut non-seed}} \geq 1$
$A_{\text{ROC}} = 0.54$
$p = 3.44 \times 10^{-2}$
F. miRNASNP-v3, mutations related to other diseases and traits
$N_{\text{mut seed}} \geq 1, N_{\text{mut non-seed}} \geq 1$
$A_{\text{ROC}} = 0.521$
True positive rate
False positive rate
random
$\Delta p_{\text{unfolded seed}} = p_{\text{unfolded seed mutant}} - p_{\text{unfolded seed WT}}$

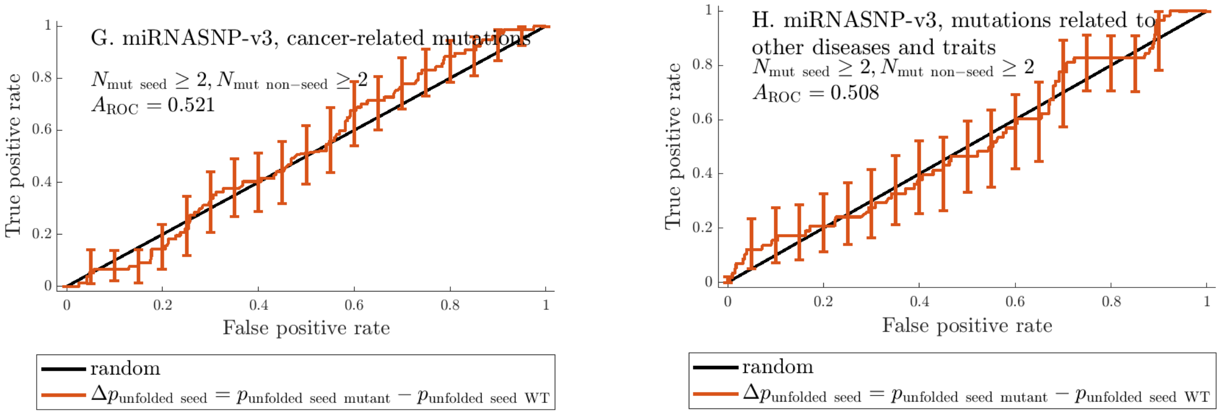

Figure 3: ROC curves built using $\Delta p_{\text{unfolded seed}} = p_{\text{unfolded seed mutant}} - p_{\text{unfolded seed WT}}$ as the criterion for predicting disease-related mutations. Data taken from SomamiR and miRNASNP-v3.

###### 2.2.2 $\langle d_{\text{Hamming}} \rangle$ percentile

Additional ROC curves for data from miRNASNP-v3 are shown in Figure 6.

#### 2.3 Positional-entropy-based criteria

##### 2.3.1 Change of the positional entropy of the mutated site with respect to the WT

We use the change in positional entropy $S^{(i)}$ at the mutated site as defined in Eq. (3) as a criterion for predicting whether a mutation is associated with disease and present the corresponding ROC curves for various datasets in Figure 8.

##### 2.3.2 Mutant average positional entropy

The average positional entropy of an RNA, $\langle \Delta S \rangle$ (as defined in Eq. (4), is a measure of the stability of its fold. We use it as a criterion for predicting whether a mutation is associated with disease and present the corresponding ROC curves for various datasets in Figure 9.

$\langle S_{\text{mutant}} \rangle$, performs significantly better than random for the set of 28 cancer-related miRNAs with $N_{\text{mut seed}} \geq 2$ and $N_{\text{mut non-seed}} \geq 2$, indicating that disease-related mutants in this set tend to have a more stable fold than other mutants.

The change in the average positional entropy with respect to the wild type, $\langle \Delta S \rangle$, performs significantly better than random for the set of 9 miRNAs associated with diseases other than cancer for which $N_{\text{mut seed}} \geq 2$ and $N_{\text{mut non-seed}} \geq 2$. The $p$-value in this case is almost an order of magnitude higher than that for the subset of 6 miRNAs which have folded mutants and WTs, and the criterion performs no better than random for the other 3 miRNAs. This indicates that the significant effect is in the change of stability of non-trivial folds, and that, as expected, it is only appropriate to compare $\langle \Delta S \rangle$ if 1) both the mutant and the WT are folded or 2) neither is folded.

##### 2.3.3 Change of the average positional entropy with respect to the WT

If the secondary structure of a microRNA affects its interaction with its targets, then one may hypothesize that the mutations which significantly change the stability of that structure would be the ones with the

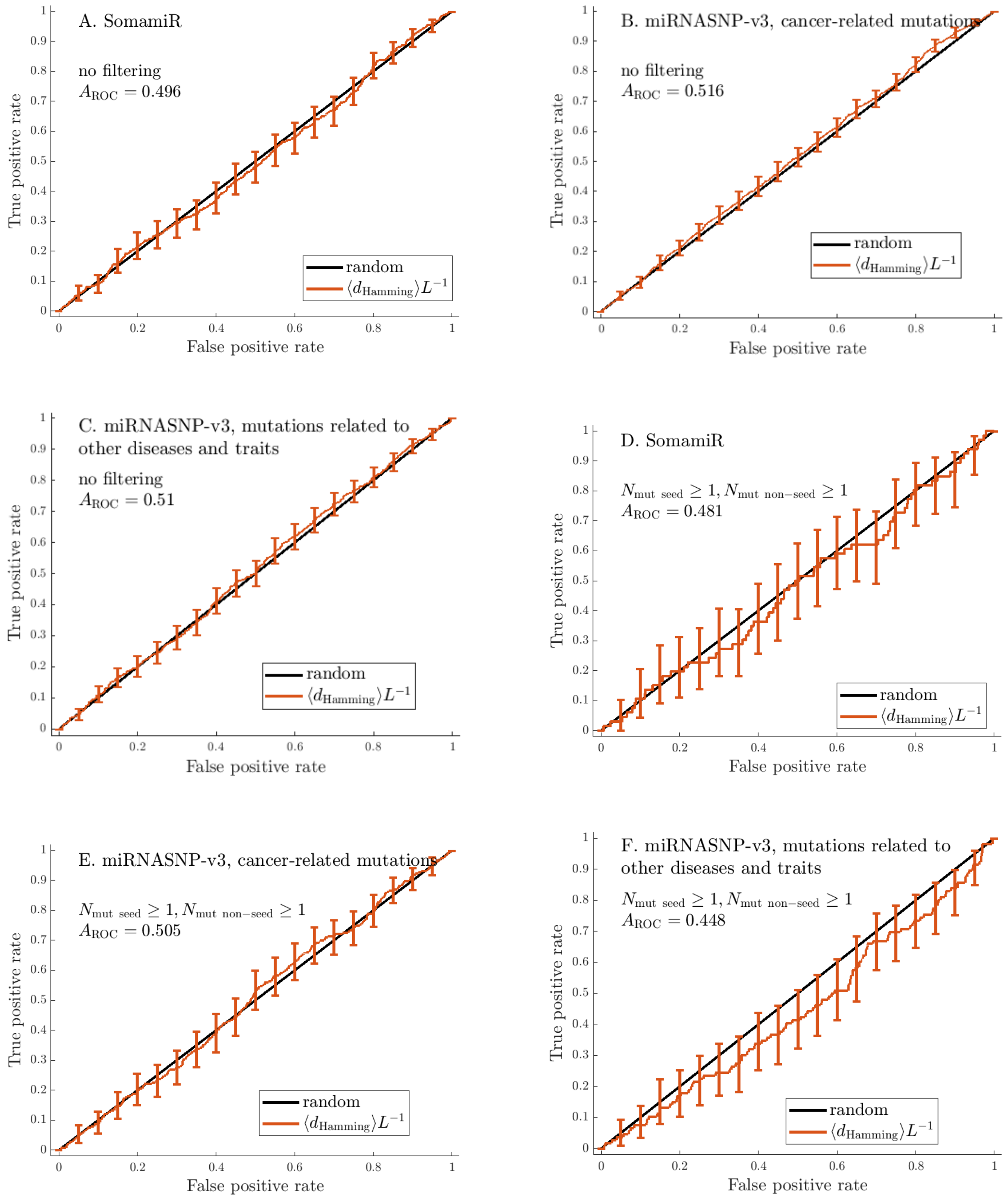
A. SomamiR
no filtering
$A_{\text{ROC}} = 0.496$
B. miRNASNP-v3, cancer-related mutations
no filtering
$A_{\text{ROC}} = 0.516$
C. miRNASNP-v3, mutations related to other diseases and traits
no filtering
$A_{\text{ROC}} = 0.51$
D. SomamiR
$N_{\text{mut seed}} \geq 1, N_{\text{mut non-seed}} \geq 1$
$A_{\text{ROC}} = 0.481$
E. miRNASNP-v3, cancer-related mutations
$N_{\text{mut seed}} \geq 1, N_{\text{mut non-seed}} \geq 1$
$A_{\text{ROC}} = 0.505$
F. miRNASNP-v3, mutations related to other diseases and traits
$N_{\text{mut seed}} \geq 1, N_{\text{mut non-seed}} \geq 1$
$A_{\text{ROC}} = 0.448$
random
$\langle d_{\text{Hamming}} \rangle L^{-1}$
True positive rate
False positive rate
0
0.1
0.2
0.3
0.4
0.5
0.6
0.7
0.8
0.9
1

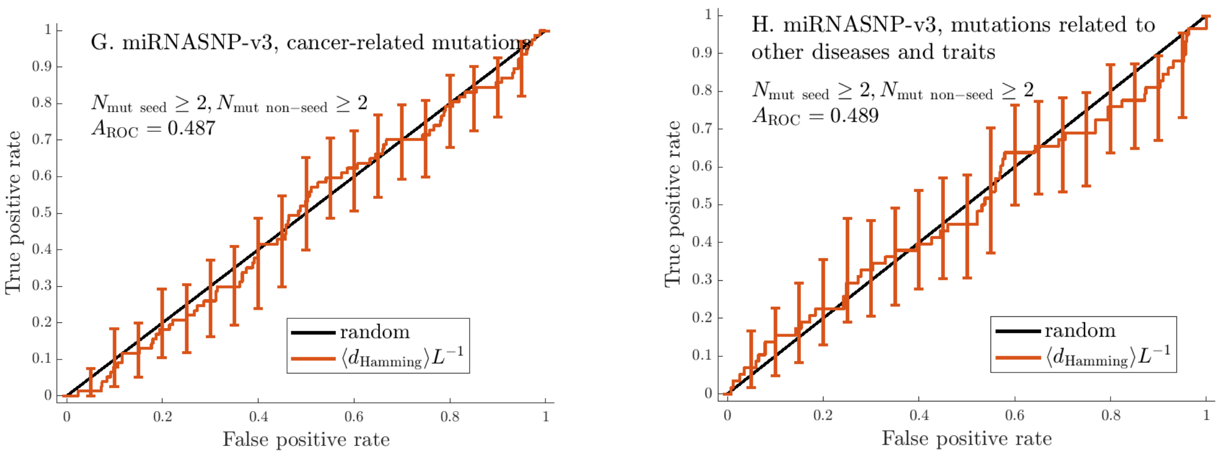

Figure 4: ROC curves built using $\langle d_{\text{Hamming}} \rangle L^{-1}$ as the criterion for predicting disease-related mutations. Data taken from SomamiR and miRNASNP-v3. The labels indicate the areas under the curves and the $p$-values for the criteria that perform significantly better than the random predictor (based on the two-sided Mann-Whitney test).

greatest effect on miRNA function. We use the difference between the average positional entropy for the mutant and the WT, $\langle \Delta S \rangle = \langle S_{\text{mut}} \rangle - \langle S_{\text{WT}} \rangle$ as a measure of this change in stability. We rank mutants according to their value of this quantity and check whether it correlates with disease association; we plot ROC curves for this criterion in Figure 10.

Strictly, the values of $\langle S \rangle$ for the mutant and the WT are comparable only if 1) the minimum-free-energy (MFE) structures both of them have secondary structure or 2) neither MFE structure has secondary structure. We analyze these two cases separately below.

**2.3.3.0.1 Folded mutant and WT** When only miRNAs with $N_{\text{mut seed}} \geq 2$ and $N_{\text{mut non-seed}} \geq 2$ from miRNASNP-v3 data are considered, the change in the average positional entropy $\langle \Delta S \rangle$ becomes a useful predictor of disease association for mutants and WT that are both folded, see Figure 11. The values of $A_{\text{ROC}}$ of 0.580 and 0.615 that we calculated for mutations related to cancer and other diseases respectively, indicate that decreasing stability is associated with disease for the clusters of 18 and 9 miRNAs in these subsets of data.

A miRNA's folding may have an effect on its function in two ways - it could either have its own functional purpose, or it could affect the interaction with the target site as it would make the binding sites within the miRNA less accessible. In the first case, a mutant may disrupt function by causing a change to the MFE secondary structure or making it less stable; in the second one, it a mutation may interfere with miRNA function by causing sites essential to target binding to enter base pairs. Our metrics aim to quantify different types of secondary structure changes in order to detect any kind of association between modified miRNA secondary structure and disease.

**2.3.3.0.2 Unfolded mutant and WT** As miRNAs need to bind to mRNAs to perform their function, one may expect that the fully unfolded state with no base pairs is ideal for performing their functions. Under this hypothesis, mutations that make the unfolded state less stable would disrupt miRNA function. The ROC curves for this subset of the data are illustrated in Figure 13. As one can see in Tables 1-8, it is much rarer for both the mutant and the WT to be unfolded and therefore, the sample size is much smaller than for the case with folded mutant and WT. This, combined with the many effects unrelated to miRNA

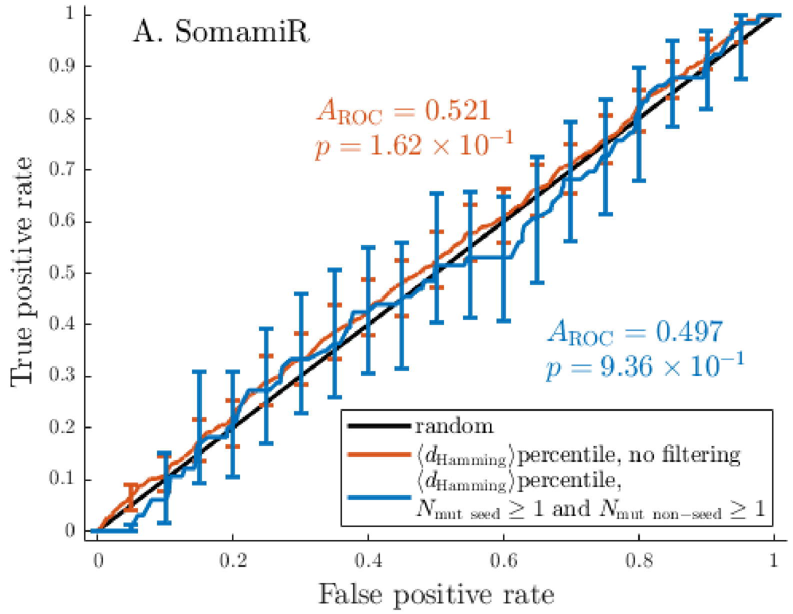
A. SomamiR
$A_{\mathrm{ROC}} = 0.521$
$p = 1.62 \times 10^{-1}$
$A_{\mathrm{ROC}} = 0.497$
$p = 9.36 \times 10^{-1}$
random
$\langle d_{\mathrm{Hamming}} \rangle$percentile, no filtering
$\langle d_{\mathrm{Hamming}} \rangle$percentile, $N_{\mathrm{mut\ seed}} \geq 1$ and $N_{\mathrm{mut\ non-seed}} \geq 1$
True positive rate
False positive rate

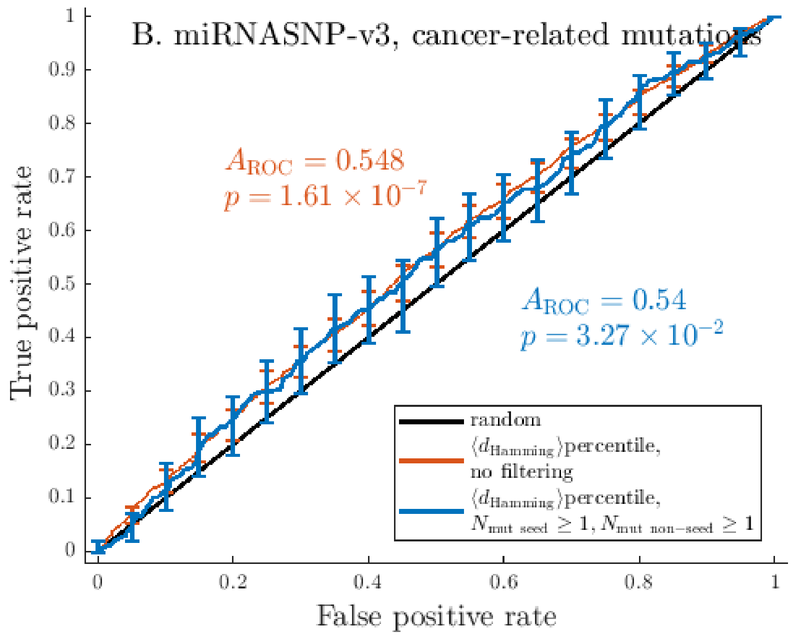
B. miRNASNP-v3, cancer-related mutations
$A_{\mathrm{ROC}} = 0.548$
$p = 1.61 \times 10^{-7}$
$A_{\mathrm{ROC}} = 0.54$
$p = 3.27 \times 10^{-2}$
random
$\langle d_{\mathrm{Hamming}} \rangle$percentile, no filtering
$\langle d_{\mathrm{Hamming}} \rangle$percentile, $N_{\mathrm{mut\ seed}} \geq 1, N_{\mathrm{mut\ non-seed}} \geq 1$
True positive rate
False positive rate

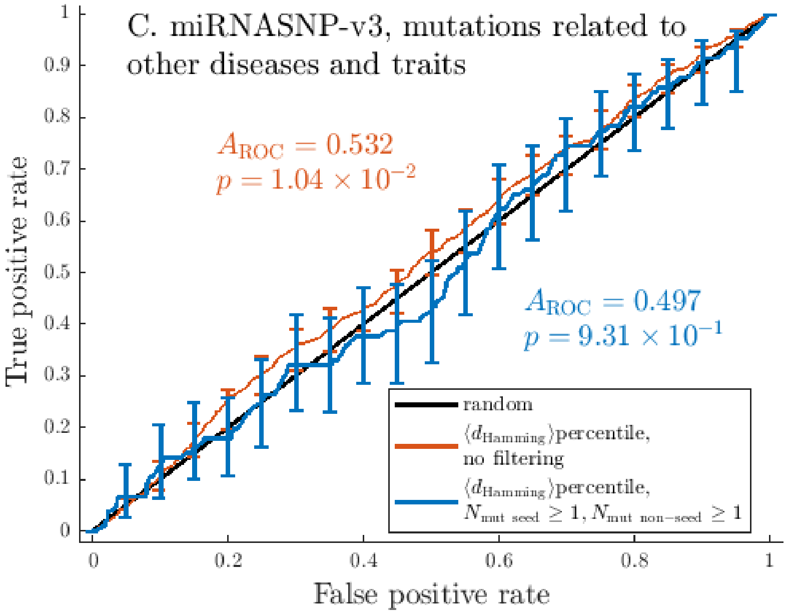

Figure 5: **The change in SS associated with a mutation as measured by the normalized Hamming distance in SS between the WT and mutated ensembles is a predictor of the mutation's relationship with disease.** ROC curves built using $\Delta p_{\mathrm{unfolded}} = p_{\mathrm{unfolded\ mutant}} - p_{\mathrm{unfolded\ WT}}$ as the criterion for predicting disease-related mutations. Data from SomamiR (A), cancer-related mutations (B) and mutations related to other traits and diseases (C) from miRNASNP-v3, with either no filtering based on $N_{\mathrm{mut\ seed}}$ and $N_{\mathrm{mut\ non-seed}}$ (red curves) or with $N_{\mathrm{mut\ seed}} \geq 1$ and $N_{\mathrm{mut\ non-seed}} \geq 1$ (blue curves) for all mature miRNAs considered here. Error bars indicate pointwise standard deviations calculated at 21 equally spaced points with the bootstrapping method [2]. The area under the ROC curve ($A_{\mathrm{ROC}}$) and the Mann-Whitney $p$-value [3] for the curves are also indicated. The $\Delta p_{\mathrm{unfolded}}$ criterion performs significantly better than the random one for both datasets, indicating that the probability that disease-related mutants are fully unfolded tends to be higher than that for other mutants. This could be because mutants with a higher $p_{\mathrm{unfolded\ mutant}}$ have a higher activity than their respective WTs, and, in the case of cancer-associated mutations, they may be more effective at downregulating tumour suppressor genes.

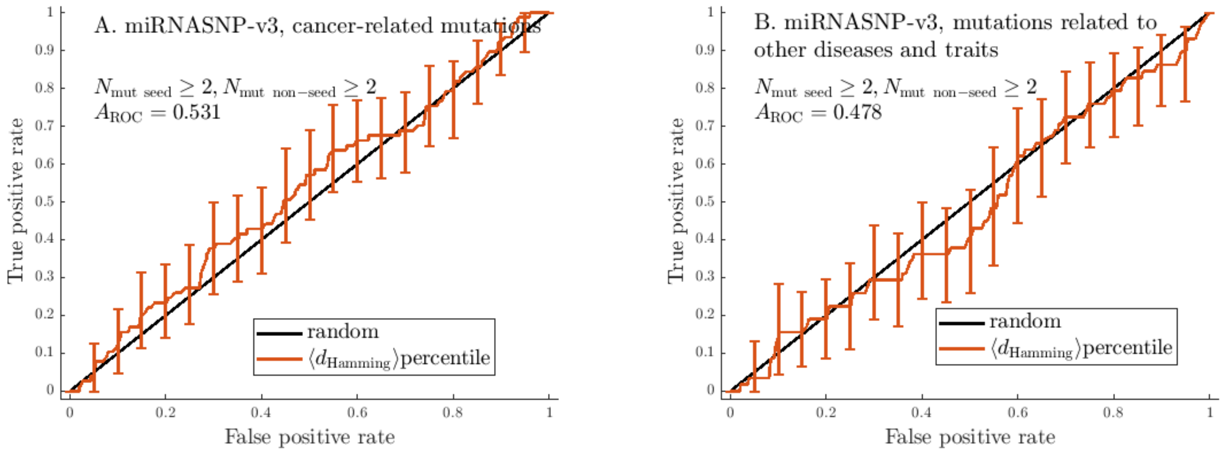

Figure 6: ROC curves built using $\langle d_{\text{Hamming}} \rangle L^{-1}$ as the criterion for predicting disease-related mutations. Data taken from miRNASNP-v3.

SS, is probably the reason that the change in entropy does not perform significantly better than random for unfolded mutant and miRNA WTs.

### 3 Tables and graphical comparisons of measures of criteria performance

### 4 Tables of $p$-values and other data for individual miRNAs

### 5 Table of disease-associated mutations that convert one WT miRNA to another

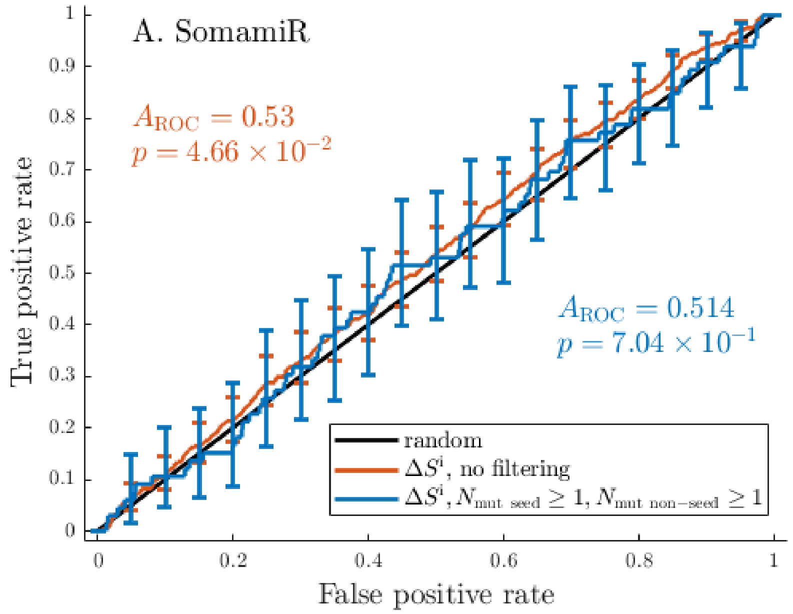

A. SomamiR
$A_{\mathrm{ROC}} = 0.53$
$p = 4.66 \times 10^{-2}$
$A_{\mathrm{ROC}} = 0.514$
$p = 7.04 \times 10^{-1}$
random
$\Delta S^{\mathrm{i}}$, no filtering
$\Delta S^{\mathrm{i}}, N_{\mathrm{mut\ seed}} \geq 1, N_{\mathrm{mut\ non-seed}} \geq 1$
True positive rate
False positive rate

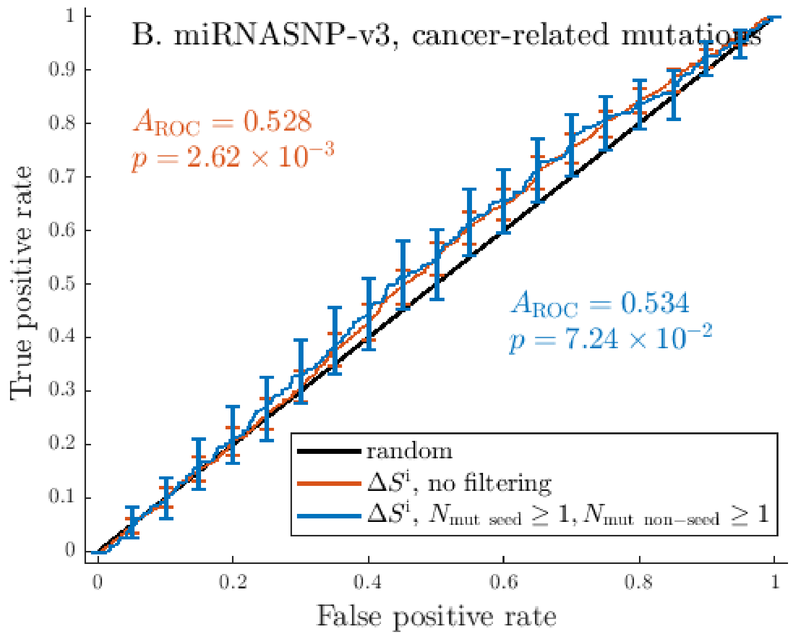

B. miRNASNP-v3, cancer-related mutations
$A_{\mathrm{ROC}} = 0.528$
$p = 2.62 \times 10^{-3}$
$A_{\mathrm{ROC}} = 0.534$
$p = 7.24 \times 10^{-2}$
random
$\Delta S^{\mathrm{i}}$, no filtering
$\Delta S^{\mathrm{i}}, N_{\mathrm{mut\ seed}} \geq 1, N_{\mathrm{mut\ non-seed}} \geq 1$
True positive rate
False positive rate

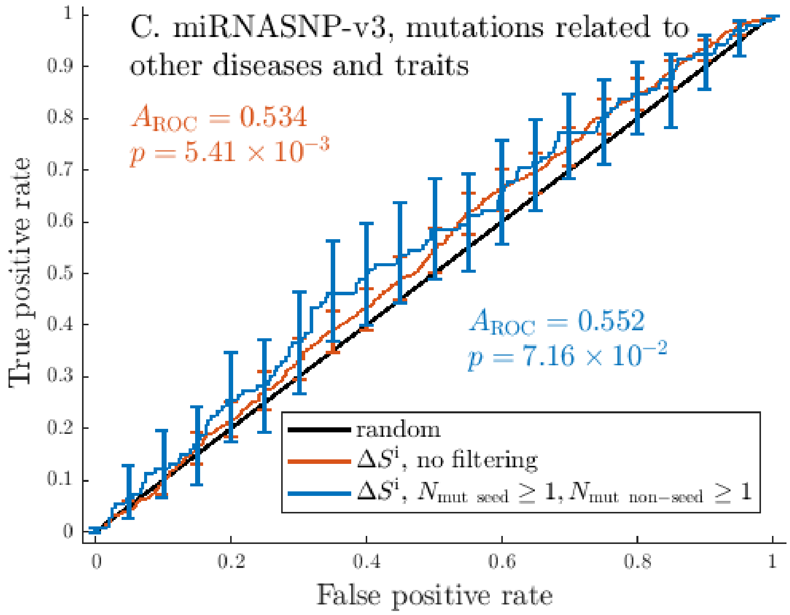

Figure 7: **The change in the positional entropy at the mutated site in a point mutant is a predictor of the mutation's relationship with disease.** ROC curves built using $\Delta S^{\mathrm{i}} = S^{\mathrm{i}}_{\mathrm{mut}} - S^{\mathrm{i}}_{\mathrm{WT}}$ as the criterion for predicting disease-related mutations. Data from SomamiR (A), cancer-related mutations (B) and mutations related to other traits and diseases (C) from miRNASNP-v3, with either no filtering based on $N_{\mathrm{mut\ seed}}$ and $N_{\mathrm{mut\ non-seed}}$ (red curves) or with $N_{\mathrm{mut\ seed}} \geq 1$ and $N_{\mathrm{mut\ non-seed}} \geq 1$ (blue curves) for all mature miRNAs considered here. Error bars indicate pointwise standard deviations calculated at 21 equally spaced points with the bootstrapping method [2]. The area under the ROC curve ($A_{\mathrm{ROC}}$) and the Mann-Whitney $p$-value [3] for the curves are also indicated. The $\Delta S^{\mathrm{i}}$ criterion performs significantly better than the random one for both datasets, when no filtering by $N_{\mathrm{mut\ seed}}$ and $N_{\mathrm{mut\ non-seed}}$ is applied. This indicates that the change in positional entropy at the mutated site tends to be lower for disease-related mutants.

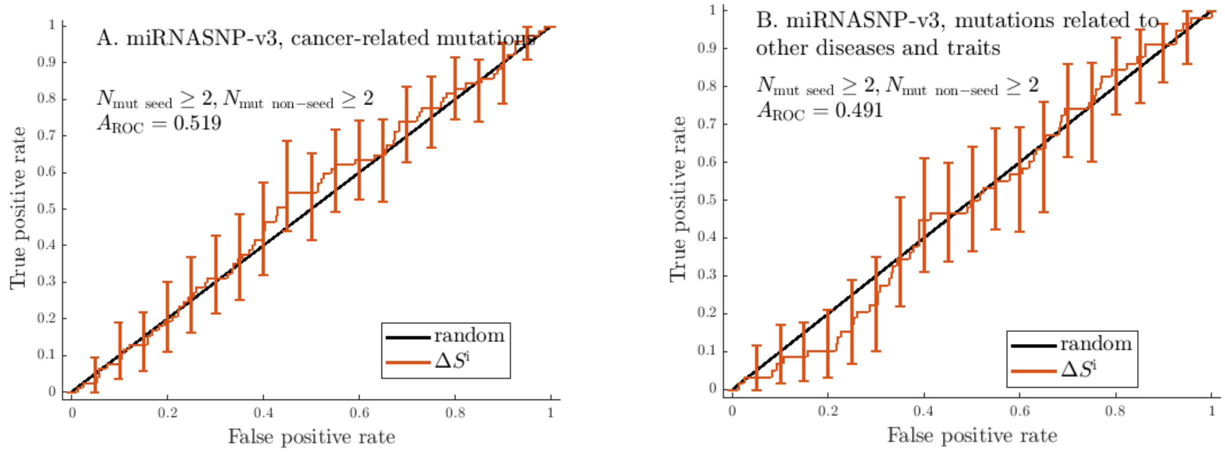

Figure 8: ROC curves built using $\langle S^{\text{i}} \rangle$ as the criterion for predicting disease-related mutations for data from SomamiR and miRNASNP-v3. The labels indicate the areas under the curves and the $p$-values for the criteria that perform significantly better than the random predictor (based on the two-sided Mann-Whitney test).

Table 1: Areas under the ROC curves ($A_{\text{ROC}}$) for various criteria applied to the miRNA entries from SomamiR with no filtering based on $N_{\text{mut seed}}$ and $N_{\text{mut non-seed}}$, and $p$-values based on the two-sided Mann-Whitney test. $p$-values of less than 0.05, which indicate criteria that perform significantly better than the random predictor (with $A_{\text{ROC}} = 0.5$), in **bold**.

| Criterion | $A_{\text{ROC}}$ | $p$-value | Number of unique mature miRNAs in the sample |
|---|---|---|---|
| $\Delta p_{\text{unfolded}}$ | 0.533 | $\mathbf{2.93 \times 10^{-2}}$ | 285 |
| $\Delta p_{\text{unfolded seed}}$ | 0.497 | $8.68 \times 10^{-1}$ | 285 |
| $\Delta S^{(\text{i})}$ | 0.530 | $\mathbf{4.66 \times 10^{-2}}$ | 285 |
| $\langle S_{\text{mut}} \rangle$ | 0.512 | $4.32 \times 10^{-1}$ | 285 |
| $\langle \Delta S \rangle$, all cases | 0.487 | $4.11 \times 10^{-1}$ | 285 |
| $\langle \Delta S \rangle$, folded mutant and WT | 0.499 | $9.51 \times 10^{-1}$ | 215 |
| $\langle \Delta S \rangle$, unfolded mutant and WT | 0.460 | $2.90 \times 10^{-1}$ | 70 |
| $\langle d_{\text{Hamming}} \rangle L^{-1}$ | 0.496 | $8.00 \times 10^{-1}$ | 285 |
| $\langle d_{\text{Hamming}} \rangle$ percentile | 0.521 | $1.62 \times 10^{-1}$ | 285 |

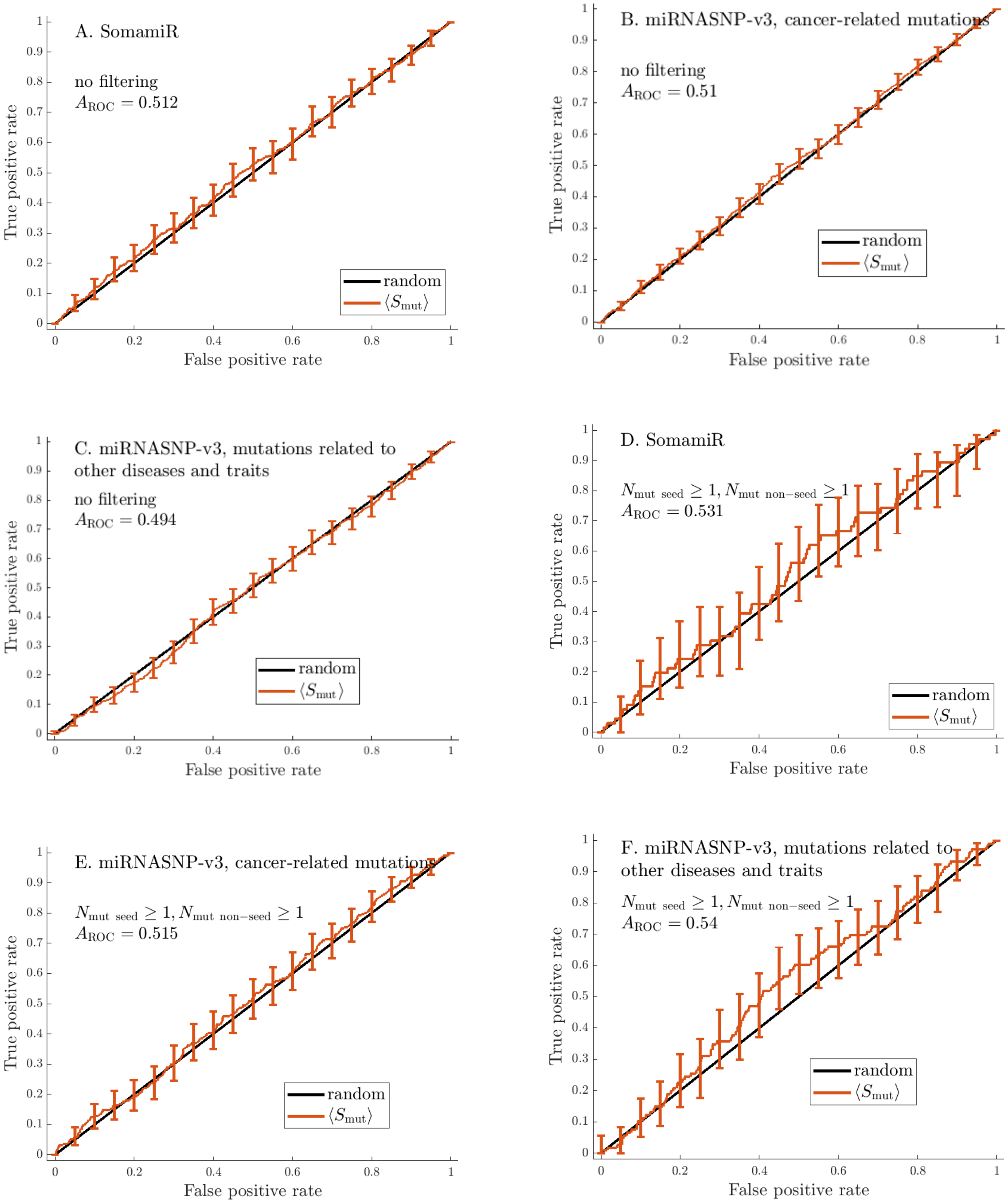
A. SomamiR
no filtering
$A_{\mathrm{ROC}} = 0.512$
B. miRNASNP-v3, cancer-related mutations
no filtering
$A_{\mathrm{ROC}} = 0.51$
C. miRNASNP-v3, mutations related to other diseases and traits
no filtering
$A_{\mathrm{ROC}} = 0.494$
D. SomamiR
$N_{\mathrm{mut\ seed}} \geq 1, N_{\mathrm{mut\ non-seed}} \geq 1$
$A_{\mathrm{ROC}} = 0.531$
E. miRNASNP-v3, cancer-related mutations
$N_{\mathrm{mut\ seed}} \geq 1, N_{\mathrm{mut\ non-seed}} \geq 1$
$A_{\mathrm{ROC}} = 0.515$
F. miRNASNP-v3, mutations related to other diseases and traits
$N_{\mathrm{mut\ seed}} \geq 1, N_{\mathrm{mut\ non-seed}} \geq 1$
$A_{\mathrm{ROC}} = 0.54$
random
$\langle S_{\mathrm{mut}} \rangle$
True positive rate
False positive rate
0
0.1
0.2
0.3
0.4
0.5
0.6
0.7
0.8
0.9
1

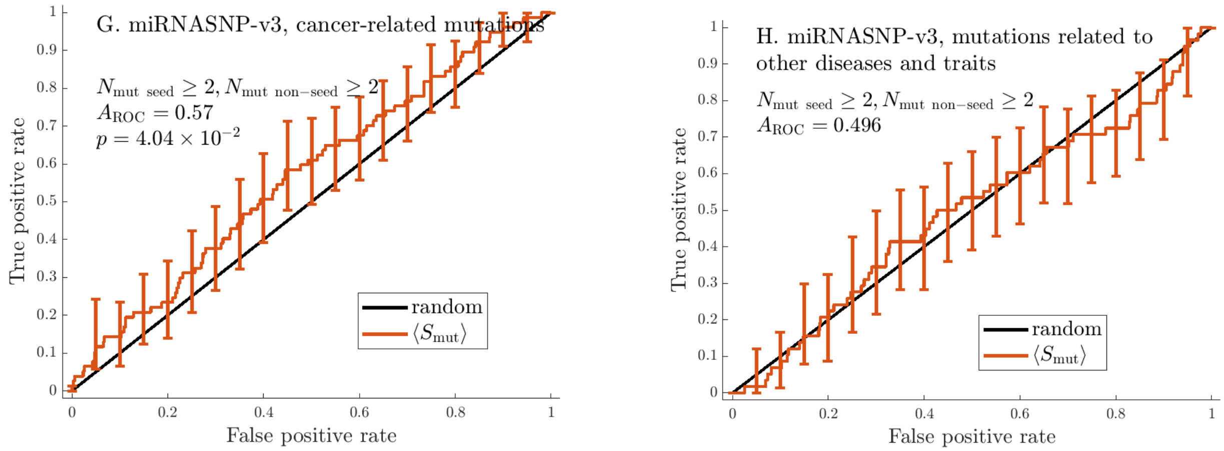

Figure 9: ROC curves built using $\langle S_{\text{mut}} \rangle$ as the criterion for predicting disease-related mutations for data from SomamiR and miRNASNP-v3. The labels indicate the areas under the curves and the $p$-values for the criteria that perform significantly better than the random predictor (based on the two-sided Mann-Whitney test).

Table 2: Results for various criteria applied to the entries from miRNASNP-v3 related to cancer; no filtering by $N_{\text{mut seed}}$ and $N_{\text{mut non-seed}}$ is applied. The table contains areas under the ROC curves ($A_{\text{ROC}}$), and $p$-values based on the two-sided Mann-Whitney test. $p$-values of less than 0.05, which indicate criteria that perform significantly better than the random predictor (with $A_{\text{ROC}} = 0.5$), in **bold**.

| Criterion | $A_{\text{ROC}}$ | $p$-value | Number of unique mature miRNAs in the sample |
|---|---|---|---|
| $\Delta p_{\text{unfolded}}$ | 0.542 | $\mathbf{5.47 \times 10^{-6}}$ | 675 |
| $\Delta p_{\text{unfolded seed}}$ | 0.504 | $6.34 \times 10^{-1}$ | 675 |
| $\Delta S^{(\text{i})}$ | 0.528 | $\mathbf{2.62 \times 10^{-3}}$ | 675 |
| $\langle S_{\text{mut}} \rangle$ | 0.510 | $2.57 \times 10^{-1}$ | 675 |
| $\langle \Delta S \rangle$, all cases | 0.492 | $3.90 \times 10^{-1}$ | 675 |
| $\langle \Delta S \rangle$, folded mutant and WT | 0.495 | $6.76 \times 10^{-1}$ | 512 |
| $\langle \Delta S \rangle$, unfolded mutant and WT | 0.474 | $2.24 \times 10^{-1}$ | 163 |
| $\langle d_{\text{Hamming}} \rangle L^{-1}$ | 0.516 | $7.83 \times 10^{-2}$ | 675 |
| $\langle d_{\text{Hamming}} \rangle$ percentile | 0.548 | $\mathbf{1.61 \times 10^{-7}}$ | 675 |

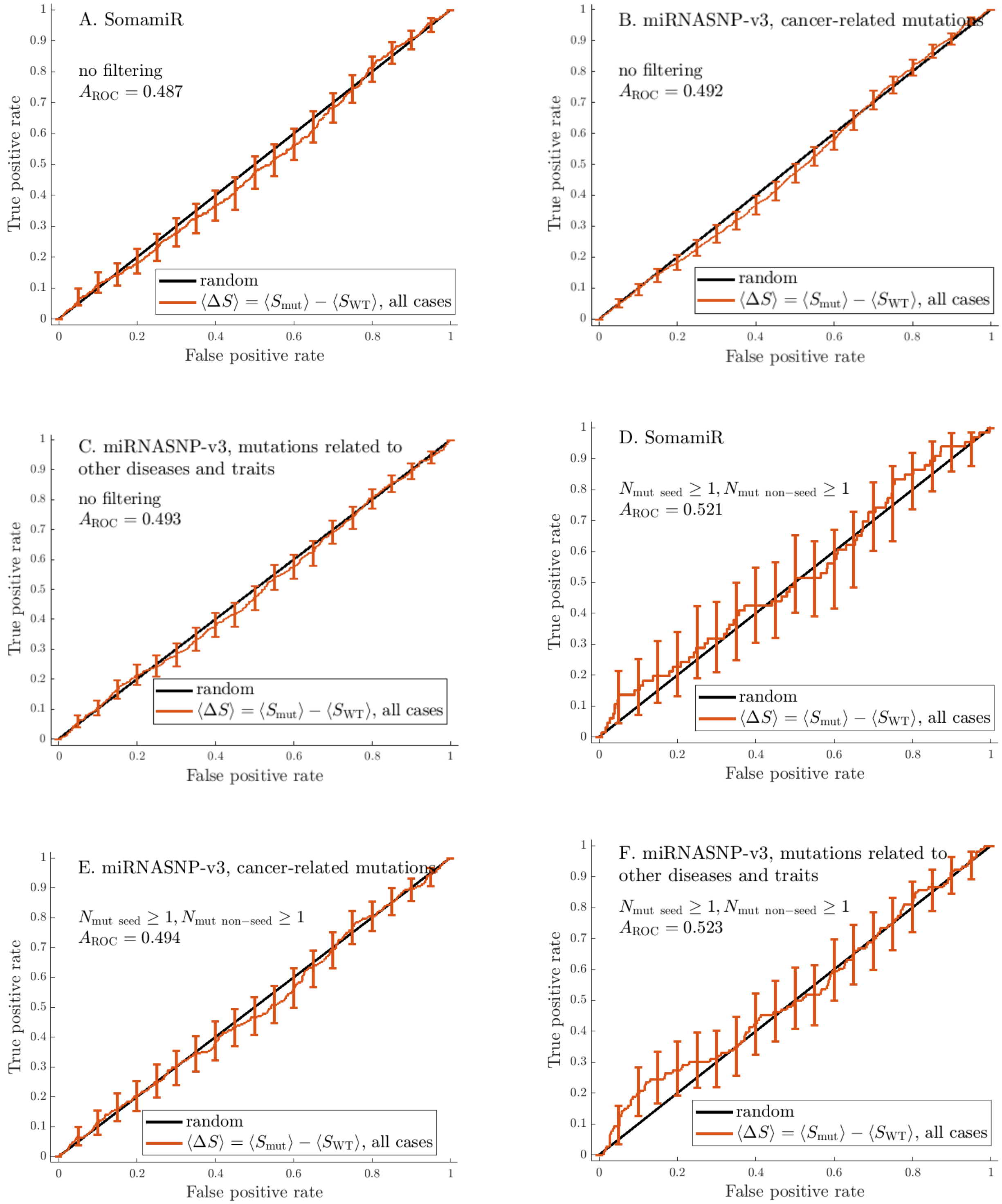
A. SomamiR
no filtering
$A_{\mathrm{ROC}} = 0.487$
B. miRNASNP-v3, cancer-related mutations
no filtering
$A_{\mathrm{ROC}} = 0.492$
C. miRNASNP-v3, mutations related to other diseases and traits
no filtering
$A_{\mathrm{ROC}} = 0.493$
D. SomamiR
$N_{\mathrm{mut\ seed}} \geq 1, N_{\mathrm{mut\ non-seed}} \geq 1$
$A_{\mathrm{ROC}} = 0.521$
E. miRNASNP-v3, cancer-related mutations
$N_{\mathrm{mut\ seed}} \geq 1, N_{\mathrm{mut\ non-seed}} \geq 1$
$A_{\mathrm{ROC}} = 0.494$
F. miRNASNP-v3, mutations related to other diseases and traits
$N_{\mathrm{mut\ seed}} \geq 1, N_{\mathrm{mut\ non-seed}} \geq 1$
$A_{\mathrm{ROC}} = 0.523$
random
$\langle \Delta S \rangle = \langle S_{\mathrm{mut}} \rangle - \langle S_{\mathrm{WT}} \rangle$, all cases
True positive rate
False positive rate
0
0.1
0.2
0.3
0.4
0.5
0.6
0.7
0.8
0.9
1

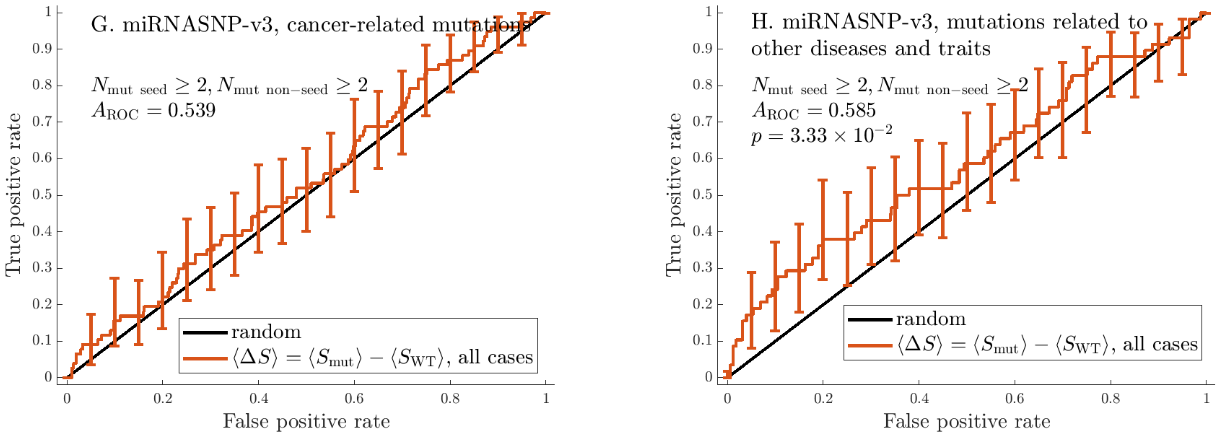

Figure 10: ROC curves built using $\langle \Delta S \rangle = \langle S_{\mathrm{mut}} \rangle - \langle S_{\mathrm{WT}} \rangle$ as the criterion for predicting disease-related mutations applied to data from SomamiR and miRNASNP-v3. The labels indicate the areas under the curves and the $p$-values for the criteria that perform significantly better than the random predictor (based on the two-sided Mann-Whitney test).

Table 3: Results for various criteria applied to the entries from miRNASNP-v3 related to diseases other than cancer; no filtering by $N_{\mathrm{mut\ seed}}$ and $N_{\mathrm{mut\ non-seed}}$ is applied. The table contains areas under the ROC curves ($A_{\mathrm{ROC}}$), and $p$-values based on the two-sided Mann-Whitney test. $p$-values of less than 0.05, which indicate criteria that perform significantly better than the random predictor (with $A_{\mathrm{ROC}} = 0.5$), in **bold**.

| Criterion | $A_{\mathrm{ROC}}$ | Number of unique mature miRNAs in the sample | |
|---|---|---|---|
| $\Delta p_{\mathrm{unfolded}}$ | 0.537 | $\mathbf{2.46 \times 10^{-3}}$ | 400 |
| $\Delta p_{\mathrm{unfolded\ seed}}$ | 0.510 | $4.28 \times 10^{-1}$ | 400 |
| $\Delta S^{(\mathrm{i})}$ | 0.534 | $\mathbf{5.41 \times 10^{-3}}$ | 400 |
| $\langle S_{\mathrm{mut}} \rangle$ | 0.494 | $6.33 \times 10^{-1}$ | 400 |
| $\langle \Delta S \rangle$, all cases | 0.493 | $5.47 \times 10^{-1}$ | 400 |
| $\langle \Delta S \rangle$, folded mutant and WT | 0.490 | $4.75 \times 10^{-1}$ | 292 |
| $\langle \Delta S \rangle$, unfolded mutant and WT | 0.459 | $1.36 \times 10^{-1}$ | 108 |
| $\langle d_{\mathrm{Hamming}} \rangle L^{-1}$ | 0.510 | $4.21 \times 10^{-1}$ | 400 |
| $\langle d_{\mathrm{Hamming}} \rangle$ percentile | 0.532 | $\mathbf{1.04 \times 10^{-2}}$ | 400 |

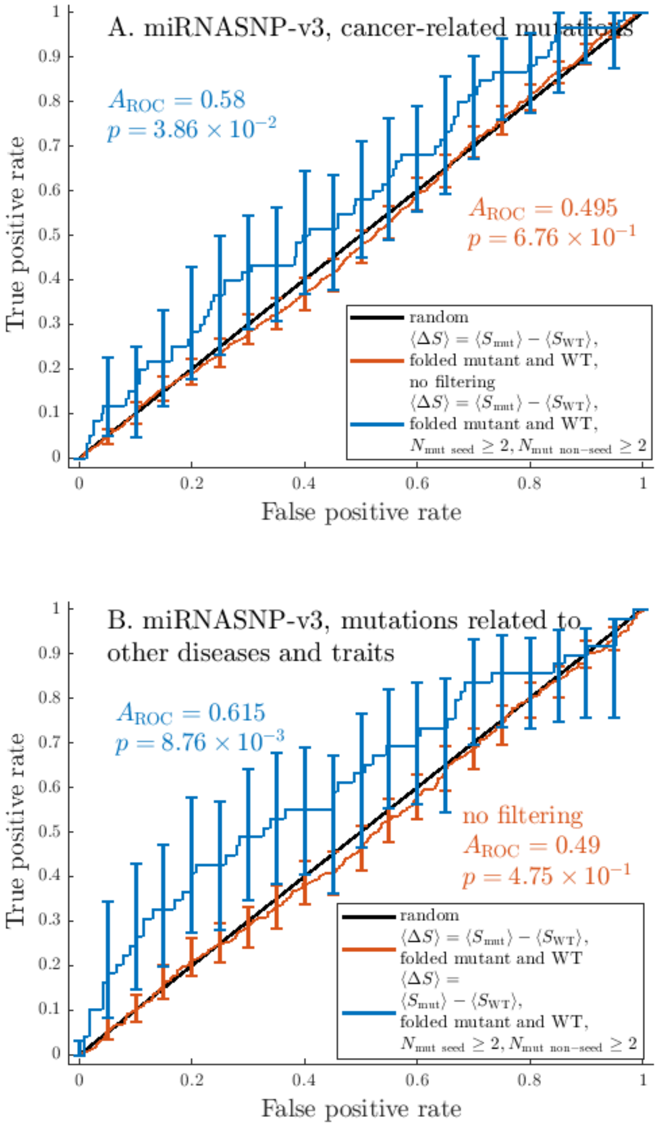

Figure 11: **The change in the average positional entropy, which measures the stability of a fold, of a miRNA associated with a mutation is a predictor of the mutation's relationship with disease for a subset of miRNAs with a comparatively large number of recorded mutations.** ROC curves built characterizing the difference in average positional entropy between mutant and WT that are both folded, $\langle \Delta S \rangle = \langle S_{\rm mut} \rangle - \langle S_{\rm WT} \rangle$. When applied to the unfiltered datasets on mutations related to cancer and other diseases from miRNASNP-v3, this criterion does not perform significantly better than the random one. However, when looking only at miRNAs for which the database records at least two mutations in both the seed and the rest of the mature region, the criterion performs significantly better than random, particularly for mutations unrelated to cancer. The results for cancer-related mutations highlight a cluster of 23 miRNAs for which the change in the stability of the fold plays a significant role, and the mutations related to other traits and diseases highlight another cluster of 9 miRNAs (see Tables 7-8 in the SI for more information on the samples). The top-ranked mutations are those with the highest $\langle \Delta S \rangle$, which suggests that for miRNAs within these clusters, less stable folds tend to be associated with disease. We explore the data for one of the miRNAs from the non-cancer cluster, hsa-miR-4537, in Figure 17, and find strong association of additional criteria with disease.

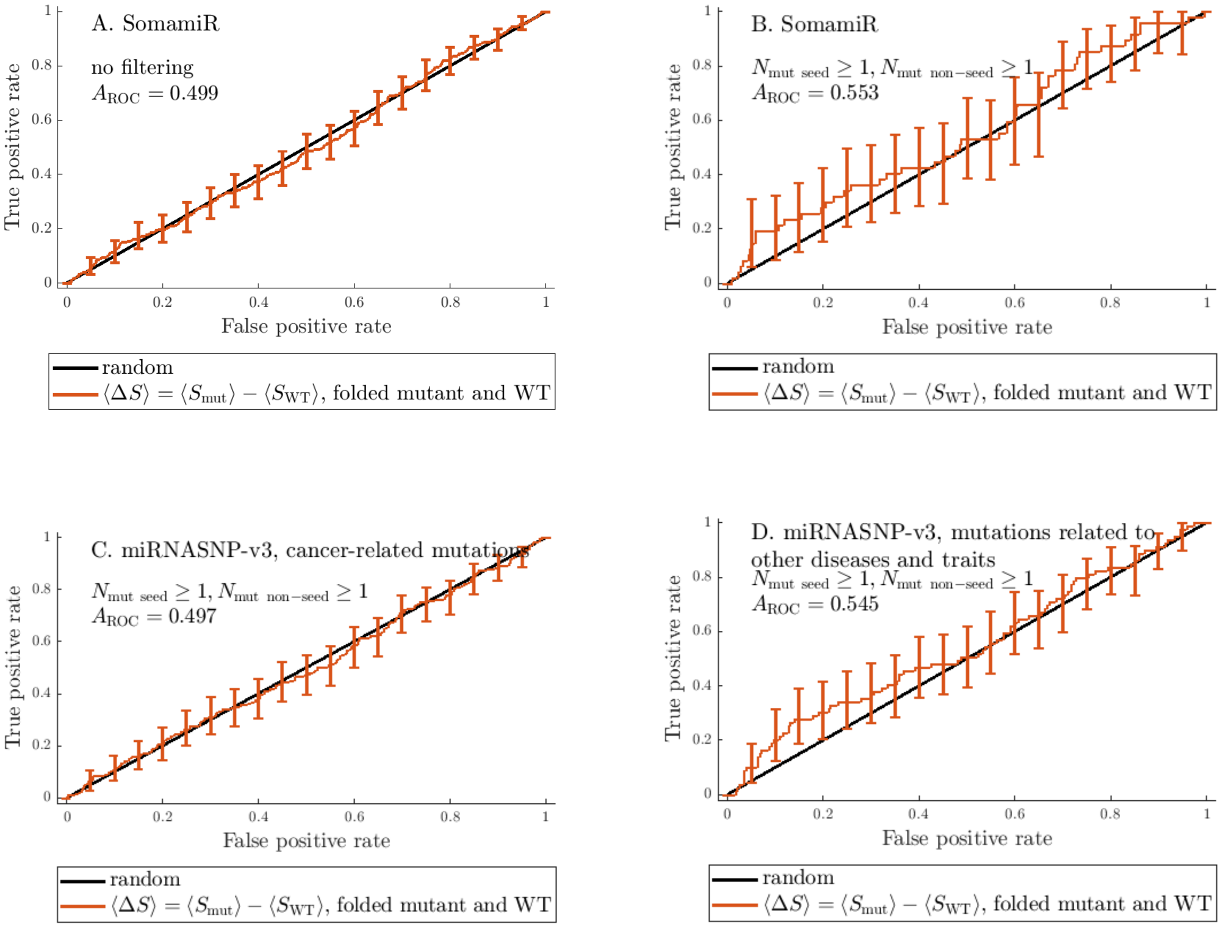

Figure 12: ROC curves built using $\langle \Delta S \rangle = \langle S_{\mathrm{mut}} \rangle - \langle S_{\mathrm{WT}} \rangle$ as the criterion for predicting disease-related mutations; the plot is based on the cases for which both the mutant and the WT MFE structures are folded. Data taken from SomamiR and miRNASNP-v3.

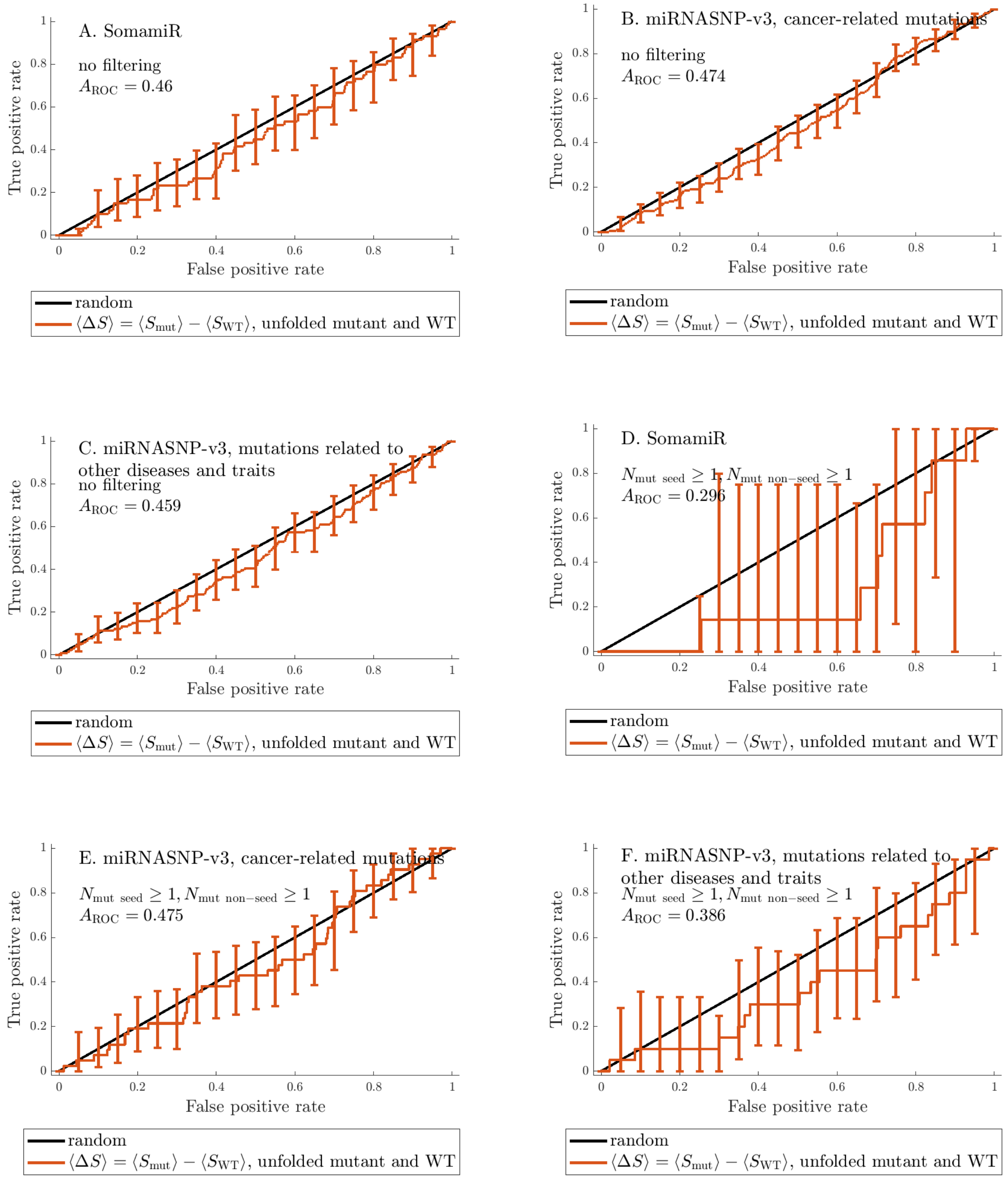
A. SomamiR
no filtering
$A_{\mathrm{ROC}} = 0.46$
B. miRNASNP-v3, cancer-related mutations
no filtering
$A_{\mathrm{ROC}} = 0.474$
C. miRNASNP-v3, mutations related to other diseases and traits
no filtering
$A_{\mathrm{ROC}} = 0.459$
D. SomamiR
$N_{\mathrm{mut\ seed}} \geq 1, N_{\mathrm{mut\ non-seed}} \geq 1$
$A_{\mathrm{ROC}} = 0.296$
E. miRNASNP-v3, cancer-related mutations
$N_{\mathrm{mut\ seed}} \geq 1, N_{\mathrm{mut\ non-seed}} \geq 1$
$A_{\mathrm{ROC}} = 0.475$
F. miRNASNP-v3, mutations related to other diseases and traits
$N_{\mathrm{mut\ seed}} \geq 1, N_{\mathrm{mut\ non-seed}} \geq 1$
$A_{\mathrm{ROC}} = 0.386$
True positive rate
False positive rate
0
0.2
0.4
0.6
0.8
1
random
$\langle \Delta S \rangle = \langle S_{\mathrm{mut}} \rangle - \langle S_{\mathrm{WT}} \rangle$, unfolded mutant and WT

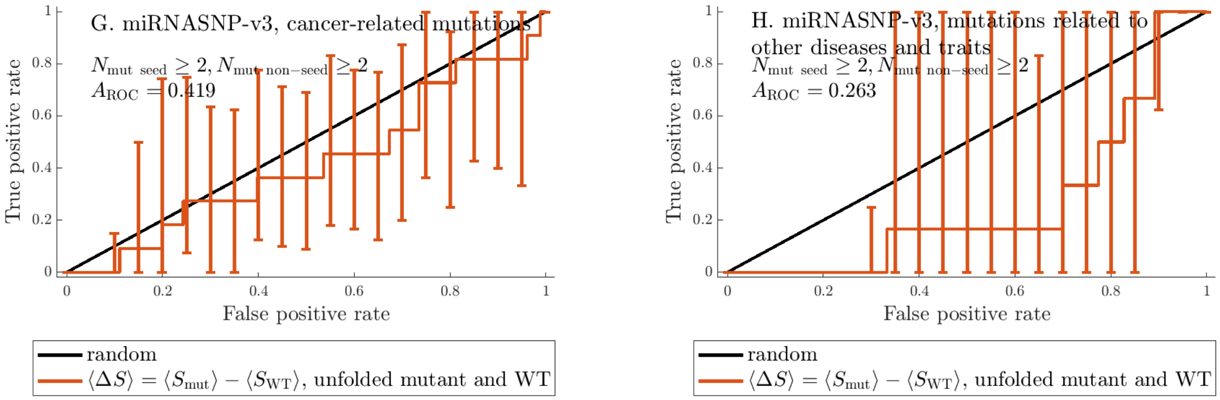

Figure 13: ROC curves built using $\langle \Delta S \rangle = \langle S_{\text{mut}} \rangle - \langle S_{\text{WT}} \rangle$ as the criterion for predicting disease-related mutations; the plot is based on the cases for which both the mutant and the WT MFE structures are fully unfolded. Data taken from SomamiR and miRNASNP-v3.

Table 4: Areas under the ROC curves ($A_{\text{ROC}}$) for various criteria applied to the miRNA entries from SomamiR for which $N_{\text{mut seed}} \geq 1$ and $N_{\text{mut non-seed}} \geq 1$, and $p$-values based on the two-sided Mann-Whitney test. $p$-values of less than 0.05, which indicate criteria that perform significantly better than the random predictor (with $A_{\text{ROC}} = 0.5$), in **bold**.

| Criterion | $A_{\text{ROC}}$ | $p$-value | Number of unique mature miRNAs in the sample |
|---|---|---|---|
| $\Delta p_{\text{unfolded}}$ | 0.579 | $\mathbf{2.88 \times 10^{-2}}$ | 40 |
| $\Delta p_{\text{unfolded seed}}$ | 0.540 | $2.70 \times 10^{-1}$ | 40 |
| $\Delta S^{(\text{i})}$ | 0.514 | $7.04 \times 10^{-1}$ | 40 |
| $\langle S_{\text{mut}} \rangle$ | 0.531 | $3.93 \times 10^{-1}$ | 40 |
| $\langle \Delta S \rangle$, all cases | 0.521 | $5.64 \times 10^{-1}$ | 40 |
| $\langle \Delta S \rangle$, folded mutant and WT | 0.553 | $2.16 \times 10^{-1}$ | 31 |
| $\langle \Delta S \rangle$, unfolded mutant and WT | 0.296 | $6.59 \times 10^{-2}$ | 9 |
| $\langle d_{\text{Hamming}} \rangle L^{-1}$ | 0.481 | $5.96 \times 10^{-1}$ | 40 |
| $\langle d_{\text{Hamming}} \rangle$ percentile | 0.497 | $9.36 \times 10^{-1}$ | 40 |

A. Performance of various criteria when applied to SomamiR data

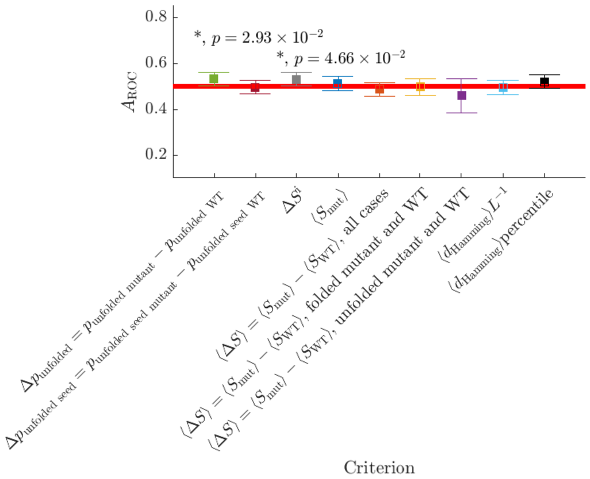

B. Performance of various criteria when applied to miRNASNP-v3 data, cancer-related mutations

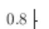

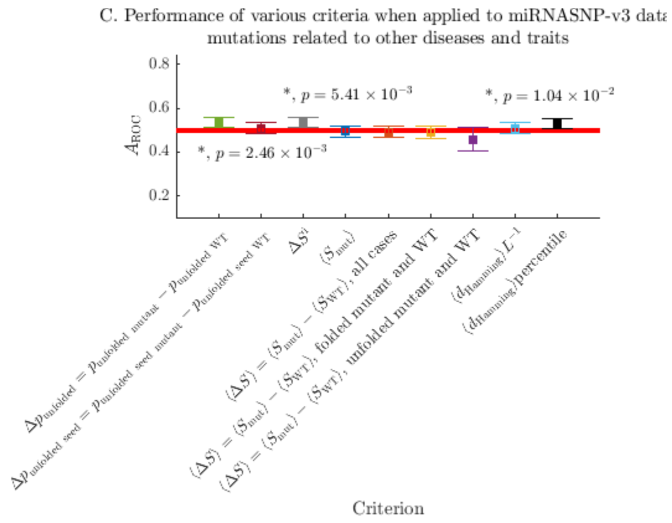

Figure 14: **Two independent SS-based criteria consistently predict association of miRNA mutations with disease better than the random criterion.** Comparison of the performance of various criteria in terms of predicting disease-associated mutations when applied to data from SomamiR (A), data on cancer-related mutations (B) and mutations related to other traits and diseases (C) from miRNASNP-v3 with no filtering based on $N_{\mathrm{mut\ seed}}$ and $N_{\mathrm{mut\ non-seed}}$. Squares mark $A_{\mathrm{ROC}}$ values, error bars show 95% confidence intervals, and Mann-Whitney $p$-values are indicated wherever $p < 0.05$. The red horizontal lines indicate the area under the curve for the random criterion, $A_{\mathrm{ROC}} = 0.5$. The criterion that performs significantly better than random for all datasets is $\Delta p_{\mathrm{unfolded}}$, which measures the change in probability that the miRNA is fully unfolded. For miRNASNP-v3 data, $A_{\mathrm{ROC}}$ is also significantly greater 0.5 if mutants are instead ranked in ascending order of their percentile in the $\langle d_{\mathrm{Hamming}} \rangle$ distribution. This means that, within the two miRNASNP-v3 datasets, mutations which change miRNA secondary structure less tend to be associated with disease.

A. Performance of various criteria when applied to SomamiR data

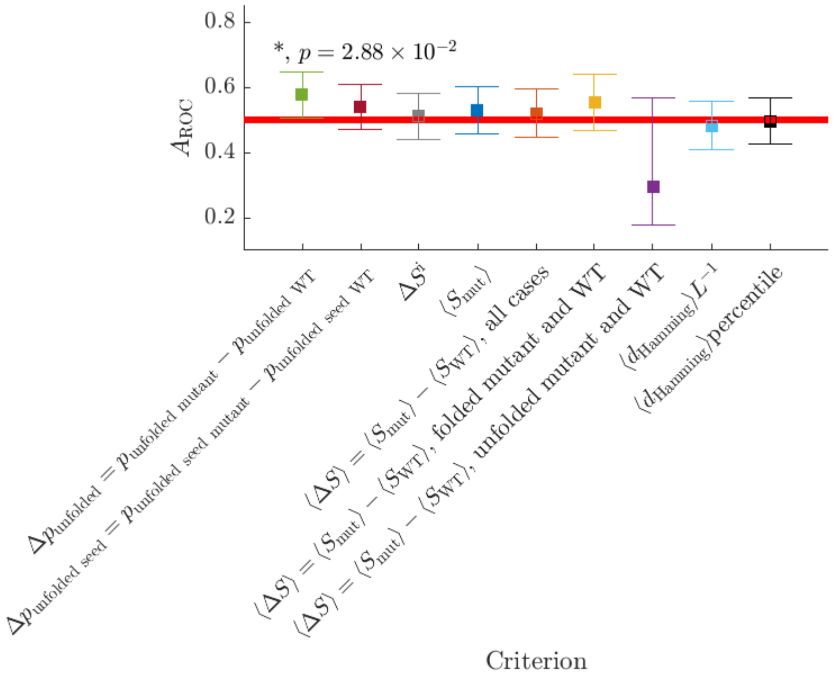

B. Performance of various criteria when applied to miRNASNP-v3 dat
cancer-related mutations

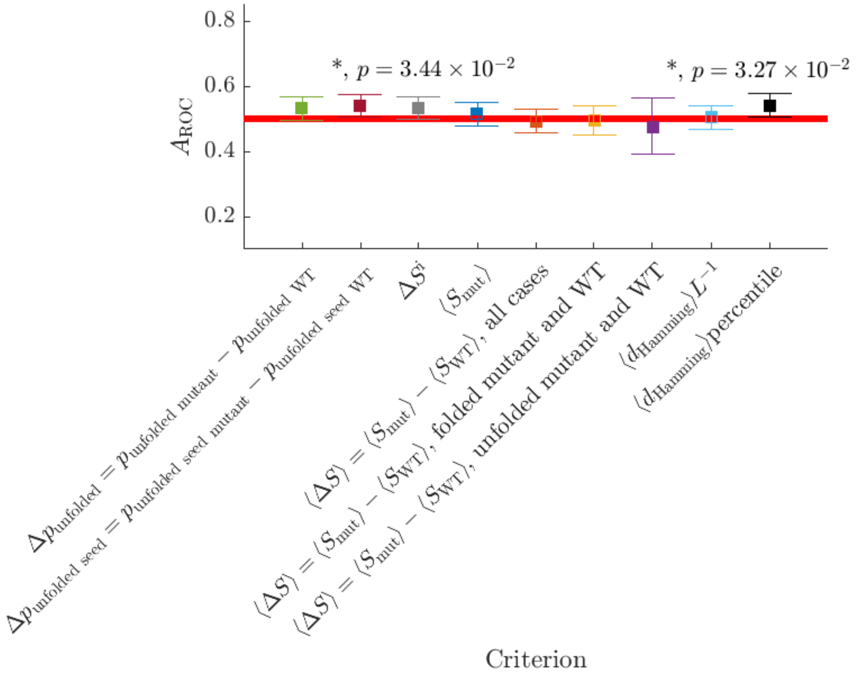

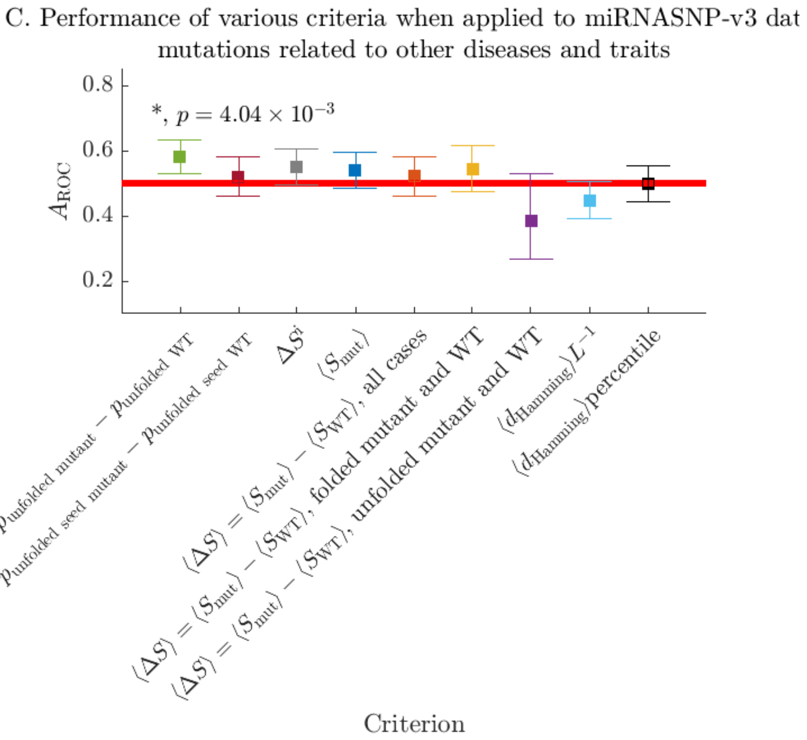

Figure 15: Comparison of the performance of various criteria in terms of predicting disease-associated mutations when applied to data from SomamiR and miRNASNP-v3 with $N_{\mathrm{mut\ seed}} \geq 1$ and $N_{\mathrm{mut\ non-seed}} \geq 1$. Squares mark $A_{\mathrm{ROC}}$ values, error bars show 95% confidence intervals, and $p$-values are indicated wherever $p < 0.05$. The red horizontal line indicates the area under the curve for the random criterion, $A_{\mathrm{ROC}} = 0.5$.

Table 5: Results for various criteria applied to the entries from miRNASNP-v3 related to cancer for which $N_{\mathrm{mut\ seed}} \geq 1$ and $N_{\mathrm{mut\ non-seed}} \geq 1$. The table contains areas under the ROC curves ($A_{\mathrm{ROC}}$), and $p$-values based on the two-sided Mann-Whitney test. $p$-values of less than 0.05, which indicate criteria that perform significantly better than the random predictor (with $A_{\mathrm{ROC}} = 0.5$), in **bold**.

| Criterion | $A_{\mathrm{ROC}}$ | $p$-value | Number of unique mature miRNAs in the sample |
|---|---|---|---|
| $\Delta p_{\mathrm{unfolded}}$ | 0.533 | $7.94 \times 10^{-2}$ | 143 |
| $\Delta p_{\mathrm{unfolded\ seed}}$ | 0.540 | $\mathbf{3.44 \times 10^{-2}}$ | 143 |
| $\Delta S^{(\mathrm{i})}$ | 0.534 | $7.24 \times 10^{-2}$ | 143 |
| $\langle S_{\mathrm{mut}} \rangle$ | 0.515 | $4.18 \times 10^{-1}$ | 143 |
| $\langle \Delta S \rangle$, all cases | 0.494 | $7.42 \times 10^{-1}$ | 143 |
| $\langle \Delta S \rangle$, folded mutant and WT | 0.497 | $8.85 \times 10^{-1}$ | 109 |
| $\langle \Delta S \rangle$, unfolded mutant and WT | 0.475 | $5.85 \times 10^{-1}$ | 34 |
| $\langle d_{\mathrm{Hamming}} \rangle L^{-1}$ | 0.505 | $8.09 \times 10^{-1}$ | 143 |
| $\langle d_{\mathrm{Hamming}} \rangle$ percentile | 0.540 | $\mathbf{3.27 \times 10^{-2}}$ | 143 |

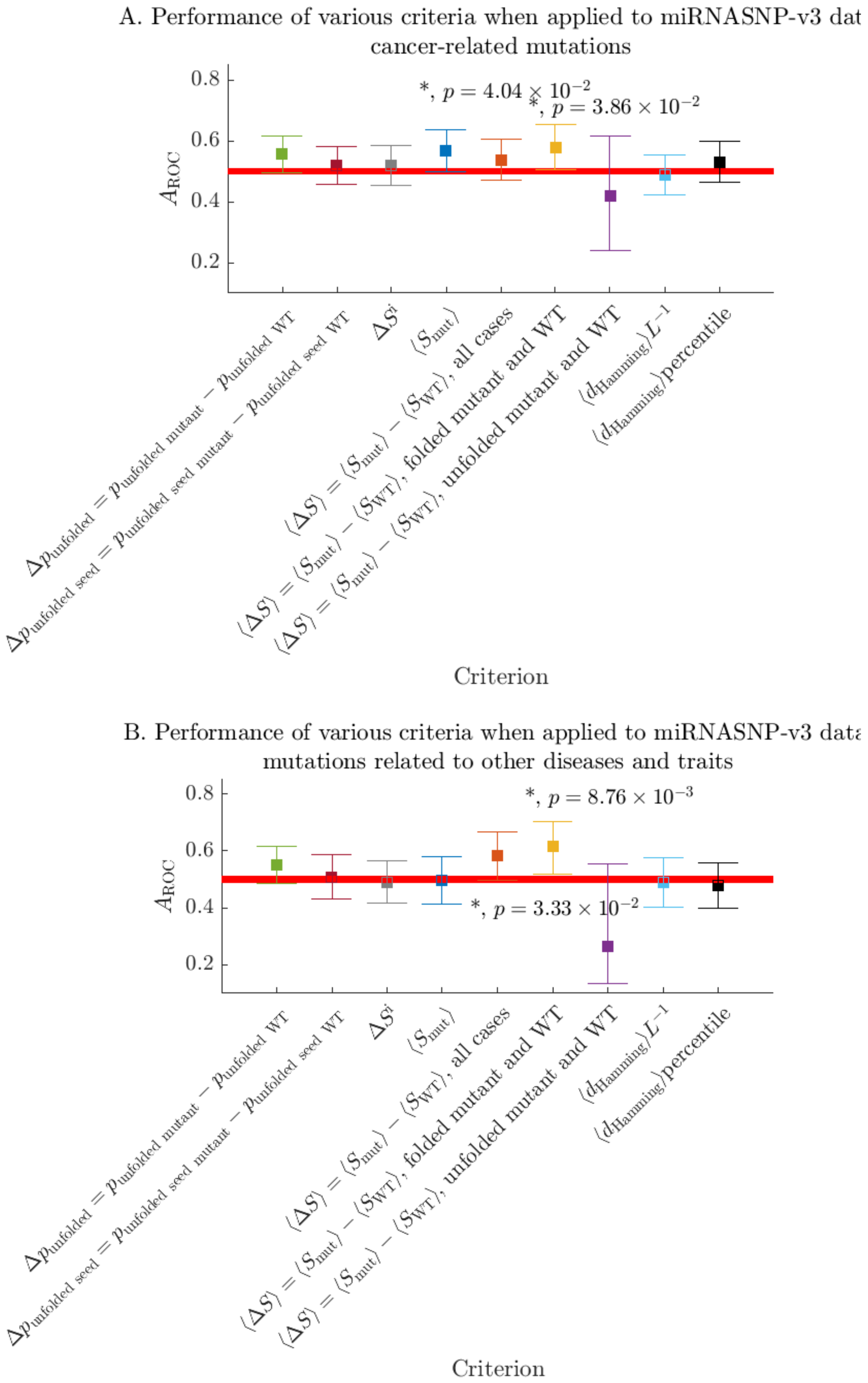

Figure 16: Comparison of the performance of various criteria in terms of predicting disease-associated mutations when applied to data from miRNASNP-v3 with $N_{\mathrm{mut\ seed}} \geq 2$ and $N_{\mathrm{mut\ non-seed}} \geq 2$. Squares mark $A_{\mathrm{ROC}}$ values, error bars show 95% confidence intervals, and $p$-values are indicated wherever $p < 0.05$. The red horizontal line indicates the area under the curve for the random criterion, $A_{\mathrm{ROC}} = 0.5$.

Table 6: Results for various criteria applied to the entries from miRNASNP-v3 related to diseases other than cancer for which $N_{\text{mut seed}} \geq 1$ and $N_{\text{mut non-seed}} \geq 1$. The table contains areas under the ROC curves ($A_{\text{ROC}}$), and $p$-values based on the two-sided Mann-Whitney test. $p$-values of less than 0.05, which indicate criteria that perform significantly better than the random predictor (with $A_{\text{ROC}} = 0.5$), in **bold**.

| Criterion | $A_{\text{ROC}}$ | $p$-value | Number of unique mature miRNAs in the sample |
|---|---|---|---|
| $\Delta p_{\text{unfolded}}$ | 0.583 | $\mathbf{4.04 \times 10^{-3}}$ | 46 |
| $\Delta p_{\text{unfolded seed}}$ | 0.521 | $4.70 \times 10^{-1}$ | 46 |
| $\Delta S^{(\text{i})}$ | 0.552 | $7.16 \times 10^{-2}$ | 46 |
| $\langle S_{\text{mut}} \rangle$ | 0.540 | $1.63 \times 10^{-2}$ | 46 |
| $\langle \Delta S \rangle$, all cases | 0.523 | $4.27 \times 10^{-1}$ | 46 |
| $\langle \Delta S \rangle$, folded mutant and WT | 0.545 | $1.75 \times 10^{-1}$ | 32 |
| $\langle \Delta S \rangle$, unfolded mutant and WT | 0.386 | $8.31 \times 10^{-2}$ | 14 |
| $\langle d_{\text{Hamming}} \rangle L^{-1}$ | 0.448 | $7.17 \times 10^{-2}$ | 46 |
| $\langle d_{\text{Hamming}} \rangle$ percentile | 0.497 | $9.31 \times 10^{-1}$ | 46 |

Table 7: Results for various criteria applied to the entries from miRNASNP-v3 related to cancer for which $N_{\text{mut seed}} \geq 2$ and $N_{\text{mut non-seed}} \geq 2$. The table contains areas under the ROC curves ($A_{\text{ROC}}$), and $p$-values based on the two-sided Mann-Whitney test. $p$-values of less than 0.05, which indicate criteria that perform significantly better than the random predictor (with $A_{\text{ROC}} = 0.5$), in **bold**.

| Criterion | $A_{\text{ROC}}$ | $p$-value (> 95% confidence) | Number of unique mature miRNAs in the sample |
|---|---|---|---|
| $\Delta p_{\text{unfolded}}$ | 0.557 | $9.26 \times 10^{-2}$ | 23 |
| $\Delta p_{\text{unfolded seed}}$ | 0.521 | $5.29 \times 10^{-1}$ | 23 |
| $\Delta S^{(\text{i})}$ | 0.519 | $5.71 \times 10^{-1}$ | 23 |
| $\langle S_{\text{mut}} \rangle$ | 0.570 | $\mathbf{4.04 \times 10^{-2}}$ | 23 |
| $\langle \Delta S \rangle$, all cases | 0.539 | $2.56 \times 10^{-1}$ | 23 |
| $\langle \Delta S \rangle$, folded mutant and WT | 0.580 | $\mathbf{3.86 \times 10^{-2}}$ | 18 |
| $\langle \Delta S \rangle$, unfolded mutant and WT | 0.419 | $3.68 \times 10^{-1}$ | 5 |
| $\langle d_{\text{Hamming}} \rangle L^{-1}$ | 0.487 | $7.06 \times 10^{-1}$ | 23 |
| $\langle d_{\text{Hamming}} \rangle$ percentile | 0.531 | $3.70 \times 10^{-1}$ | 23 |

Table 8: Results for various criteria applied to the entries from miRNASNP-v3 related to diseases other than cancer for which $N_{\text{mut seed}} \geq 2$ and $N_{\text{mut non-seed}} \geq 2$. The table contains areas under the ROC curves ($A_{\text{ROC}}$), and $p$-values based on the two-sided Mann-Whitney test. $p$-values of less than 0.05, which indicate criteria that perform significantly better than the random predictor (with $A_{\text{ROC}} = 0.5$), in **bold**.

| Criterion | $A_{\text{ROC}}$ | $p$-value (> 95% confidence) | Number of unique mature miRNAs in the sample |
|---|---|---|---|
| $\Delta p_{\text{unfolded}}$ | 0.552 | $1.95 \times 10^{-1}$ | 12 |
| $\Delta p_{\text{unfolded seed}}$ | 0.508 | $8.40 \times 10^{-1}$ | 12 |
| $\Delta S^{(\text{i})}$ | 0.491 | $8.19 \times 10^{-1}$ | 12 |
| $\langle S_{\text{mut}} \rangle$ | 0.496 | $9.22 \times 10^{-1}$ | 12 |
| $\langle \Delta S \rangle$, all cases | 0.585 | $\mathbf{3.33 \times 10^{-2}}$ | 12 |
| $\langle \Delta S \rangle$, folded mutant and WT | 0.615 | $\mathbf{8.76 \times 10^{-3}}$ | 9 |
| $\langle \Delta S \rangle$, unfolded mutant and WT | 0.263 | $5.38 \times 10^{-2}$ | 3 |
| $\langle d_{\text{Hamming}} \rangle L^{-1}$ | 0.489 | $7.88 \times 10^{-1}$ | 12 |
| $\langle d_{\text{Hamming}} \rangle$ percentile | 0.478 | $5.79 \times 10^{-1}$ | 12 |

Table 9: Results for individual miRNAs from SomamiR for which $\Delta p_{\text{unfolded}}$ performs better than the random criterion ($p < 0.05$). The $p$-values are based on the two-sided Mann-Whitney test, and the SS of the WT is predicted with ViennaRNA. For each miRNA, we give lists of references connecting it to cancer and other traits and diseases.

| miRNA | $A_{\text{ROC}}$ | $p$-value | Predicted SS (WT) | Ref., cancer | Ref., other traits and diseases |
|---|---|---|---|---|---|
| hsa-miR-3689b-3p | 0.076 | $3.55 \times 10^{-2}$ | ...................... | | |
| hsa-miR-19a-3p | 0.000 | $3.92 \times 10^{-2}$ | ..((((......)))))....... | [4–6] | [7–9] |
| hsa-miR-548g-5p | 0.082 | $3.92 \times 10^{-2}$ | (((((......))))))....... | [10] | |
| hsa-miR-345-3p | 1.000 | $4.17 \times 10^{-2}$ | (((((......)))))...... | [11] | [12] |
| hsa-miR-30e-5p | 1.000 | $4.17 \times 10^{-2}$ | ........(((......))).. | [13–17] | [18–20] |
| hsa-miR-367-3p | 1.000 | $4.17 \times 10^{-2}$ | .(((((......))))).... | [21] | |

Table 10: Results for individual miRNAs from SomamiR for which $\langle d_{\text{Hamming}} \rangle L^{-1}$ performs better than the random criterion ($p < 0.05$). The $p$-values are based on the two-sided Mann-Whitney test, and the SS of the WT is predicted with ViennaRNA. For each miRNA, we give lists of references connecting it to cancer and other traits and diseases.

| miRNA | $A_{\text{ROC}}$ | $p$-value | Predicted SS (WT) | Ref., cancer | Ref., other traits and diseases |
|---|---|---|---|---|---|
| hsa-miR-3939 | 0.142 | $1.52 \times 10^{-2}$ | .....(((..(....)..))). | | [22] |
| hsa-miR-520g-3p | 1.000 | $3.70 \times 10^{-2}$ | .......((((.......)))).. | [23] | |
| hsa-miR-635 | 1.000 | $3.92 \times 10^{-2}$ | ....(((((.((...)).))))) | [24, 25] | |
| hsa-miR-1185-1-3p | 1.000 | $4.17 \times 10^{-2}$ | ......(((((....))))).. | [26] | [27] |
| hsa-miR-208a-3p | 1.000 | $4.17 \times 10^{-2}$ | .....((((((.....)))))) | | [28, 29] |
| hsa-miR-887-3p | 1.000 | $4.17 \times 10^{-2}$ | .....((((......))))... | [30, 31] | [32] |
| hsa-miR-192-3p | 1.000 | $4.17 \times 10^{-2}$ | ..(((.........)))..... | [33] | [34–36] |
| hsa-miR-1294 | 1.000 | $4.17 \times 10^{-2}$ | .........((((....)))). | [37, 38] | [39] |
| hsa-miR-125b-2-3p | 1.000 | $4.17 \times 10^{-2}$ | ((.((((.......)))).)). | [40, 41] | |
| hsa-miR-485-5p | 0.913 | $4.43 \times 10^{-2}$ | ...((....))........... | [42–54] | [55] |

Table 11: miRNAs for which the combined $p$-values from the $\Delta p_{\text{unfolded}}$ and $\langle d_{\text{Hamming}} \rangle L^{-1}$ criteria is below 0.05 based on data from SomamiR. miRNA identifiers in *italics* if the $p$-value for $\Delta p_{\text{unfolded}}$ is also less than 0.05 and **bold** if the $p$-value for $\langle d_{\text{Hamming}} \rangle L^{-1}$ is less than 0.05. As elsewhere, $p$-values are based on the two-sided Mann-Whitney test.

| miRNA | combined $p$-value | Predicted SS (WT) | Ref., cancer | Ref., other traits and diseases |
|---|---|---|---|---|
| *hsa-miR-3689b-3p* | $2.62 \times 10^{-2}$ | ...................... | | |
| **hsa-miR-1185-1-3p** | $3.26 \times 10^{-2}$ | ......(((((....))))).. | [26] | [27] |
| hsa-miR-558 | $3.29 \times 10^{-2}$ | ................... | [56] | [57] |
| hsa-miR-129-1-3p | $4.15 \times 10^{-2}$ | ...................... | [58] | [59] |
| hsa-miR-129-2-3p | $4.15 \times 10^{-2}$ | ...................... | [60] | [59, 61–63] |
| **hsa-miR-3939** | $4.16 \times 10^{-2}$ | .....(((..(....)..))). | | [22] |
| hsa-miR-543 | $4.29 \times 10^{-2}$ | ...................... | [64] | [64, 65] |
| **hsa-miR-208a-3p** | $4.99 \times 10^{-2}$ | .....((((((.....)))))) | | [28, 29] |
| **hsa-miR-887-3p** | $4.99 \times 10^{-2}$ | .....((((......))))... | [30, 31] | [32] |

Table 12: Results for individual miRNAs from miRNASNP-v3 for which $\Delta p_{\text{unfolded}}$ performs better than the random criterion ($p < 0.05$). The $p$-values are based on the two-sided Mann-Whitney test, and the SS of the WT is predicted with ViennaRNA. For each miRNA, we give lists of references connecting it to cancer and other traits and diseases. The names of the miRNAs for which the $\langle d_{\text{Hamming}} \rangle L^{-1}$ criterion also performs significantly better than the random one are in **bold**.

| miRNA | $A_{\text{ROC}}$ | $p$-value | Predicted SS (WT) | Ref., cancer | Ref., other traits and diseases |
|---|---|---|---|---|---|
| **hsa-miR-3150a-5p** | 0.000 | $1.77 \times 10^{-3}$ | ..................... | | [66] |
| hsa-miR-345-3p | 1.000 | $1.77 \times 10^{-3}$ | (((((......)))))...... | [11] | [12] |
| hsa-miR-580-5p | 0.000 | $1.77 \times 10^{-3}$ | ...(((.((....)).)))... | [67] | |
| **hsa-miR-4641** | 0.990 | $2.80 \times 10^{-3}$ | ....................... | [68] | |
| hsa-miR-519a-3p | 0.052 | $3.58 \times 10^{-3}$ | .....((((.((...)).)))) | [69] | [70, 71] |
| **hsa-miR-4537** | 0.746 | $5.17 \times 10^{-3}$ | ..(((.((((...)))).))). | [72] | |
| **hsa-miR-539-5p** | 0.933 | $6.13 \times 10^{-3}$ | (((.......))).......... | [73–76] | |
| hsa-miR-4756-3p | 0.022 | $7.09 \times 10^{-3}$ | (((....))).......... | [12] | [77–79] |
| hsa-miR-6852-5p | 0.023 | $8.08 \times 10^{-3}$ | .(((.((...)).))).... | [80] | |
| **hsa-miR-1269b** | 0.033 | $1.06 \times 10^{-2}$ | .(((......))).......... | [81–83] | |
| hsa-miR-548i | 0.864 | $1.33 \times 10^{-2}$ | ..........(((.....))). | [84] | [85] |
| hsa-miR-548l | 0.957 | $1.60 \times 10^{-2}$ | ..................... | [86] | [87, 88] |
| hsa-miR-208a-3p | 0.054 | $2.13 \times 10^{-2}$ | .....((((((.....)))))) | | [28, 29] |
| hsa-miR-6888-3p | 0.942 | $2.42 \times 10^{-2}$ | .................... | | |
| hsa-miR-4649-3p | 0.942 | $2.42 \times 10^{-2}$ | ...((((....))))...... | | [89, 90] |
| **hsa-miR-6821-3p** | 0.930 | $3.23 \times 10^{-2}$ | .................... | | [91] |
| hsa-miR-4518 | 0.860 | $3.36 \times 10^{-2}$ | .(((((.(..(....)..).))))). | [92, 93] | [94] |
| **hsa-miR-19a-5p** | 0.924 | $3.55 \times 10^{-2}$ | .((..((((....))))..)). | [95] | [96, 97] |
| hsa-miR-190b-5p | 0.076 | $3.55 \times 10^{-2}$ | .(((((.....)))))...... | [98] | [99] |
| hsa-miR-1307-3p | 0.924 | $3.55 \times 10^{-2}$ | ..((((((...)))))).... | | |
| hsa-miR-362-5p | 1.000 | $3.70 \times 10^{-2}$ | ...(((........)))....... | [100] | |
| hsa-miR-608 | 0.918 | $3.76 \times 10^{-2}$ | .(((((.((......)).))))).. | [101–107] | [108, 109] |
| hsa-miR-323b-5p | 0.854 | $3.90 \times 10^{-2}$ | .((....))((((.....)))). | | [110, 111] |
| hsa-miR-548g-5p | 0.082 | $3.92 \times 10^{-2}$ | (((((......)))))....... | [10] | |
| hsa-miR-657 | 0.000 | $3.92 \times 10^{-2}$ | ((.(((.......))).)).... | [112, 113] | |
| hsa-miR-19b-2-5p | 1.000 | $4.17 \times 10^{-2}$ | .....((((....)))).... | | [114] |
| hsa-miR-3116 | 1.000 | $4.17 \times 10^{-2}$ | ..((((........)))).... | [115] | |
| hsa-miR-30e-5p | 1.000 | $4.17 \times 10^{-2}$ | ........(((......))).. | [13–17] | [18–20] |
| hsa-miR-367-3p | 1.000 | $4.17 \times 10^{-2}$ | .(((((......))))).... | [21] | |
| hsa-miR-382-5p | 1.000 | $4.17 \times 10^{-2}$ | (((....)))............ | [116–120] | [121, 122] |
| **hsa-miR-558** | 0.081 | $4.32 \times 10^{-2}$ | ................... | [56] | [57] |
| hsa-miR-492 | 0.847 | $4.40 \times 10^{-2}$ | .(..((...))..)......... | [123–129] | |
| hsa-miR-518b | 0.087 | $4.43 \times 10^{-2}$ | ....((.(((.......))))) | | [130] |
| **hsa-miR-1307-5p** | 1.000 | $4.44 \times 10^{-2}$ | ....((((.......)))).. | | |
| **hsa-miR-6893-5p** | 1.000 | $4.44 \times 10^{-2}$ | ..................... | [131] | |
| **hsa-miR-892c-5p** | 1.000 | $4.44 \times 10^{-2}$ | ..................... | | |
| hsa-miR-520b-3p | 1.000 | $4.44 \times 10^{-2}$ | .........((((...)))). | [132] | |
| hsa-miR-8088 | 1.000 | $4.44 \times 10^{-2}$ | ((((..........)))).. | | [133] |
| hsa-miR-3120-5p | 1.000 | $4.44 \times 10^{-2}$ | ......(((((.....))))) | [134] | |
| hsa-miR-3150b-3p | 0.000 | $4.44 \times 10^{-2}$ | (((.(....).)))....... | [135] | [136–138] |
| hsa-miR-563 | 0.807 | $4.64 \times 10^{-2}$ | .((..(((....))).)). | | [139–141] |
| hsa-miR-3689b-3p | 0.156 | $4.69 \times 10^{-2}$ | ...................... | | |
| **hsa-miR-1227-3p** | 1.000 | $4.76 \times 10^{-2}$ | .................... | | [142–144] |
| **hsa-miR-6794-3p** | 0.000 | $4.76 \times 10^{-2}$ | .................... | | [145] |

Table 13: Results for individual miRNAs from miRNASNP-v3 for which $\langle d_{\mathrm{Hamming}} \rangle L^{-1}$ performs better than the random criterion ($p < 0.05$). The $p$-values are based on the two-sided Mann-Whitney test, and the SS of the WT is predicted with ViennaRNA. For each miRNA, we give lists of references connecting it to cancer and other traits and diseases. The names of the miRNAs for which the $\Delta p_{\mathrm{unfolded}}$ criterion also performs significantly better than the random one are in **bold**.

| miRNA | $A_{\mathrm{ROC}}$ | $p$-value | Predicted SS (WT) | Ref., cancer | Ref., other traits and diseases |
|---|---|---|---|---|---|
| hsa-miR-1908-3p | 0.939 | $1.26 \times 10^{-3}$ | ((((...)))).......... | | |
| hsa-miR-4477b | 1.000 | $1.77 \times 10^{-3}$ | (((((..((....))..))))) | [146] | |
| hsa-miR-4722-3p | 1.000 | $1.77 \times 10^{-3}$ | ..((((.((.......)))))) | [147] | [148] |
| **hsa-miR-1269b** | 0.000 | $1.77 \times 10^{-3}$ | .(((......))).......... | [81–83] | |
| hsa-miR-485-5p | 0.926 | $1.99 \times 10^{-3}$ | ...((....))........... | [42–54] | [55] |
| **hsa-miR-4641** | 0.981 | $5.59 \times 10^{-3}$ | ........................ | [68] | |
| **hsa-miR-3150a-5p** | 0.022 | $7.09 \times 10^{-3}$ | ...................... | | [66] |
| **hsa-miR-19a-5p** | 0.043 | $1.60 \times 10^{-2}$ | .((..((((....))))..)). | [95] | [96, 97] |
| **hsa-miR-4537** | 0.289 | $1.76 \times 10^{-2}$ | ..(((.((((...)))).))). | [72] | |
| hsa-miR-6800-3p | 0.047 | $1.82 \times 10^{-2}$ | ..........(((...))).. | | |
| hsa-miR-99a-3p | 0.946 | $2.13 \times 10^{-2}$ | ...(((((......)))))... | | |
| **hsa-miR-558** | 0.054 | $2.43 \times 10^{-2}$ | ................... | [56] | [57] |
| hsa-miR-3939 | 0.195 | $2.49 \times 10^{-2}$ | .....(((..(....)..))). | | [22] |
| hsa-miR-513c-5p | 0.935 | $2.84 \times 10^{-2}$ | .((((...)))).......... | [149, 150] | [151] |
| hsa-miR-148b-5p | 0.065 | $2.84 \times 10^{-2}$ | ....((((.........)))). | | [152, 153] |
| hsa-miR-4802-3p | 0.071 | $3.14 \times 10^{-2}$ | ....((((.((...))))))... | [154] | |
| hsa-miR-1248 | 1.000 | $3.17 \times 10^{-2}$ | .............((((...))))... | [24, 155] | [156] |
| hsa-miR-509-5p | 0.930 | $3.23 \times 10^{-2}$ | (((((....))))))....... | [157–159] | [160, 161] |
| hsa-miR-3130-3p | 0.930 | $3.23 \times 10^{-2}$ | ..(((.((((...))))))). | [162] | [163] |
| **hsa-miR-6821-3p** | 0.930 | $3.23 \times 10^{-2}$ | ..................... | | [91] |
| hsa-miR-550a-5p | 0.819 | $3.35 \times 10^{-2}$ | .........(((........))) | [164] | [165] |
| hsa-miR-887-3p | 0.924 | $3.55 \times 10^{-2}$ | .....((((......))))... | [30, 31] | [32] |
| hsa-miR-624-3p | 0.857 | $3.86 \times 10^{-2}$ | ....(((((.......))))) | [166] | |
| hsa-miR-103a-1-5p | 1.000 | $3.92 \times 10^{-2}$ | (((..(........)..)))... | | |
| hsa-miR-302a-5p | 1.000 | $3.92 \times 10^{-2}$ | ........(((....))))..... | | [167] |
| hsa-miR-635 | 1.000 | $3.92 \times 10^{-2}$ | ....(((((.((...)).))))) | [24, 25] | |
| **hsa-miR-539-5p** | 0.148 | $4.14 \times 10^{-2}$ | (((.......)))......... | [73–76] | |
| hsa-miR-125b-2-3p | 1.000 | $4.17 \times 10^{-2}$ | ((.((((.......)))).)). | [40, 41] | |
| hsa-miR-1185-1-3p | 1.000 | $4.17 \times 10^{-2}$ | ......(((((....))))).. | [26] | [27] |
| hsa-miR-3915 | 1.000 | $4.17 \times 10^{-2}$ | ...((((........))))... | [168] | |
| hsa-miR-5195-5p | 1.000 | $4.17 \times 10^{-2}$ | ..((....((....))....)) | | [169] |
| hsa-miR-5682 | 1.000 | $4.17 \times 10^{-2}$ | .....((((((.....)))))) | [170] | |
| hsa-miR-199a-3p | 0.000 | $4.17 \times 10^{-2}$ | .(((....)))........... | [171–174] | [175] |
| hsa-miR-381-5p | 0.000 | $4.17 \times 10^{-2}$ | .((((((.....)))))).... | [176–178] | |
| hsa-miR-3689a-3p | 0.193 | $4.30 \times 10^{-2}$ | (((.((.........)).))). | | |
| hsa-miR-328-5p | 0.847 | $4.40 \times 10^{-2}$ | ....((((.......)))).... | [179] | [34] |
| hsa-miR-500a-3p | 0.913 | $4.43 \times 10^{-2}$ | .....(((.....)))...... | [180, 181] | |
| hsa-miR-4708-3p | 0.913 | $4.43 \times 10^{-2}$ | .((......))........... | | [182] |
| hsa-miR-378i | 1.000 | $4.44 \times 10^{-2}$ | .(((.((((...)))))))).... | [183–185] | |
| **hsa-miR-892c-5p** | 1.000 | $4.44 \times 10^{-2}$ | ...................... | | |
| hsa-miR-3155a | 1.000 | $4.44 \times 10^{-2}$ | (((.((...))..)))..... | | [186] |
| **hsa-miR-6893-5p** | 1.000 | $4.44 \times 10^{-2}$ | ...................... | [131] | |
| hsa-miR-376c-3p | 0.000 | $4.44 \times 10^{-2}$ | ........(((...)))).... | [187–189] | |
| hsa-miR-892a | 0.000 | $4.44 \times 10^{-2}$ | ..((((((.......)))))) | [190–193] | |
| **hsa-miR-1307-5p** | 0.000 | $4.44 \times 10^{-2}$ | ....((((.......)))).. | | |
| hsa-miR-508-5p | 0.803 | $4.49 \times 10^{-2}$ | .......(((.......)))... | [194] | [195] |
| **hsa-miR-1227-3p** | 1.000 | $4.76 \times 10^{-2}$ | ..................... | | [142–144] |
| **hsa-miR-6794-3p** | 0.000 | $4.76 \times 10^{-2}$ | ..................... | | [145] |

Table 14: miRNAs for which the combined $p$-values from the $\Delta p_{\text{unfolded}}$ and $\langle d_{\text{Hamming}}\rangle L^{-1}$ criteria is below 0.05 based on data from miRNASNP-v3. miRNA identifiers in *italics* if the $p$-value for $\Delta p_{\text{unfolded}}$ is also less than 0.05, **bold** if the $p$-value for $\langle d_{\text{Hamming}}\rangle L^{-1}$ is less than 0.05, and ***bold italics*** - if both are below 0.05. As elsewhere, $p$-values are based via the two-sided Mann-Whitney test.

| miRNA | combined $p$-value | Predicted SS (WT) | Ref., cancer | Ref., other traits and diseases |
|---|---|---|---|---|
| ***hsa-miR-3150a-5p*** | $1.54\times10^{-4}$ | ...................... | | [66] |
| ***hsa-miR-4641*** | $1.89\times10^{-4}$ | ........................ | [68] | |
| ***hsa-miR-1269b*** | $2.24\times10^{-4}$ | .(((......)))......... | [81–83] | |
| ***hsa-miR-4537*** | $9.36\times10^{-4}$ | ..(((.((((...)))).))). | [72] | |
| **hsa-miR-4477b** | $1.14\times10^{-3}$ | (((((..((....))..))))) | [146] | |
| **hsa-miR-1908-3p** | $1.69\times10^{-3}$ | ((((...)))).......... | | |
| *hsa-miR-345-3p* | $1.91\times10^{-3}$ | (((((......)))))...... | [11] | [12] |
| ***hsa-miR-539-5p*** | $2.35\times10^{-3}$ | (((.......)))......... | [73–76] | |
| **hsa-miR-485-5p** | $3.84\times10^{-3}$ | ...((....)).......... | [42–54] | [55] |
| **hsa-miR-4722-3p** | $3.94\times10^{-3}$ | ..((((.((.......)))))) | [147] | [148] |
| ***hsa-miR-19a-5p*** | $4.80\times10^{-3}$ | .((..((((....))))..)). | [95] | [96, 97] |
| *hsa-miR-548l* | $6.85\times10^{-3}$ | ...................... | [86] | [87, 88] |
| ***hsa-miR-6821-3p*** | $8.22\times10^{-3}$ | ..................... | | [91] |
| ***hsa-miR-558*** | $8.25\times10^{-3}$ | ................... | [56] | [57] |
| *hsa-miR-4756-3p* | $8.81\times10^{-3}$ | (((....)))........... | [12] | [77–79] |
| *hsa-miR-6888-3p* | $1.11\times10^{-2}$ | ..................... | | |
| *hsa-miR-580-5p* | $1.14\times10^{-2}$ | ...(((.((....)).)))... | [67] | |
| ***hsa-miR-892c-5p*** | $1.43\times10^{-2}$ | ..................... | | |
| ***hsa-miR-1307-5p*** | $1.43\times10^{-2}$ | ....((((.......)))).. | | |
| ***hsa-miR-6893-5p*** | $1.43\times10^{-2}$ | ..................... | [131] | |
| ***hsa-miR-1227-3p*** | $1.61\times10^{-2}$ | .................... | | [142–144] |
| ***hsa-miR-6794-3p*** | $1.61\times10^{-2}$ | .................... | | [145] |
| **hsa-miR-6800-3p** | $1.66\times10^{-2}$ | ..........(((...))).. | | |
| hsa-miR-4434 | $2.09\times10^{-2}$ | .................. | | [196] |
| *hsa-miR-519a-3p* | $2.12\times10^{-2}$ | .....((((.((...)).)))) | [69] | [70, 71] |
| *hsa-miR-563* | $2.29\times10^{-2}$ | .((..(((....))).)). | | [139–141] |
| **hsa-miR-199a-3p** | $2.31\times10^{-2}$ | .(((....)))........... | [171–174] | [175] |
| **hsa-miR-5682** | $2.31\times10^{-2}$ | .....((((((.....)))))) | [170] | |
| hsa-miR-4296 | $2.43\times10^{-2}$ | ...(((((....))))) | [197] | [198] |
| *hsa-miR-548i* | $2.48\times10^{-2}$ | ..........(((.....))). | [84] | [85] |
| hsa-let-7f-1-3p | $2.65\times10^{-2}$ | ...................... | [199] | [200–202] |
| *hsa-miR-3689b-3p* | $2.75\times10^{-2}$ | ...................... | | |
| *hsa-miR-492* | $2.83\times10^{-2}$ | .(..((...))..)......... | [123–129] | |
| *hsa-miR-4518* | $2.87\times10^{-2}$ | .(((((.(..(....)..).))))). | [92, 93] | [94] |
| *hsa-miR-4649-3p* | $3.09\times10^{-2}$ | ...((((....))))...... | | [89, 90] |
| **hsa-miR-1185-1-3p** | $3.26\times10^{-2}$ | ......(((((....))))).. | [26] | [27] |
| **hsa-miR-4802-3p** | $3.37\times10^{-2}$ | ....((((.((...))))))... | [154] | |
| **hsa-miR-513c-5p** | $3.43\times10^{-2}$ | .((((...)))).......... | [149, 150] | [151] |
| *hsa-miR-208a-3p* | $3.61\times10^{-2}$ | .....((((((.....)))))) | | [28, 29] |
| *hsa-miR-6852-5p* | $3.77\times10^{-2}$ | .(((.((...)).))).... | [80] | |
| *hsa-miR-19b-2-5p* | $4.15\times10^{-2}$ | .....((((....)))).... | | [114] |
| hsa-miR-129-2-3p | $4.15\times10^{-2}$ | ...................... | [60] | [59, 61–63] |
| **hsa-miR-3915** | $4.15\times10^{-2}$ | ...((((........))))... | [168] | |
| hsa-miR-4441 | $4.34\times10^{-2}$ | ((((........)))). | [203] | |
| **hsa-miR-376c-3p** | $4.61\times10^{-2}$ | ........(((...))).... | [187–189] | |
| hsa-miR-520b-3p | $4.61\times10^{-2}$ | .........((((...)))). | [132] | |
| **hsa-miR-3130-3p** | $4.61\times10^{-2}$ | ..(((.((((...))))))). | [162] | [163] |
| hsa-miR-6769b-3p | $4.61\times10^{-2}$ | ..................... | [204, 205] | |
| hsa-miR-7151-3p | $4.61\times10^{-2}$ | .....((((......)))).. | | [206] |

Table 15: Data on mutations that convert one miRNA to another and are associated with disease; data from SomamiR and miRNASNP-v3.

| miRNA WT | mutated miRNA | WT sequence | mutated sequence | SS, WT | SS, mutant | $\langle d_{\text{Hamming}} \rangle \times L^{-1}$ | $\Delta p_{\text{unfolded}}$ |
|---|---|---|---|---|---|---|---|
| hsa-miR-1269a | hsa-miR-1269b | CUGGACUGAGCC$\underline{\text{A}}$UGCUACUGG | CUGGACUGAGCC$\underline{\text{G}}$UGCUACUGG | .(((......)))......... | .(((......)))......... | 0.35 | -0.0086 |
| hsa-miR-3689a-3p | hsa-miR-3689c, hsa-miR-3689b-3p | CUGGGAGGUGUGAUAU$\underline{\text{U}}$GUGGU | CUGGGAGGUGUGAUAU$\underline{\text{C}}$GUGGU | (((.((.........)).))). | ...................... | 0.34 | 0.51 |
| hsa-miR-3689b-3p | hsa-miR-3689a-3p | CUGGGAGGUGUGAUAU$\underline{\text{C}}$GUGGU | CUGGGAGGUGUGAUAU$\underline{\text{U}}$GUGGU | ...................... | (((.((.........)).))). | 0.34 | -0.51 |
| hsa-miR-3689c | has-miR-3689a-3p | CUGGGAGGUGUGAUAU$\underline{\text{C}}$GUGGU | CUGGGAGGUGUGAUAU$\underline{\text{U}}$GUGGU | ...................... | (((.((.........)).))). | 0.34 | -0.51 |
| hsa-miR-518e-5p | hsa-miR-526a-5p, hsa-miR-520c-5p, hsa-miR-518d-5p | CUCUAGAGGGAAGC$\underline{\text{A}}$CUUUCUG | CUCUAGAGGGAAGC$\underline{\text{G}}$CUUUCUG | ...(((((((.....))))))) | ...(((((((.....))))))) | 0.084 | $3.2 \times 10^{-4}$ |
| hsa-miR-519a-5p | hsa-miR-526a-5p, hsa-miR-520c-5p, hsa-miR-518d-5p | CUCUAGAGGGAAGC$\underline{\text{A}}$CUUUCUG | CUCUAGAGGGAAGC$\underline{\text{G}}$CUUUCUG | ...(((((((.....))))))) | ...(((((((.....))))))) | 0.084 | $3.2 \times 10^{-4}$ |
| hsa-miR-520c-5p | hsa-miR-519c-5p, hsa-miR-519b-5p, hsa-miR-523-5p, hsa-miR-518e-5p, hsa-miR-522-5p, hsa-miR-519a-5p | CUCUAGAGGGAAGC$\underline{\text{G}}$CUUUCUG | CUCUAGAGGGAAGC$\underline{\text{A}}$CUUUCUG | ...(((((((.....))))))) | ...(((((((.....))))))) | 0.084 | $-3.2 \times 10^{-4}$ |
| hsa-miR-522-5p | has-miR-526a-5p, hsa-miR-520c-5p, hsa-miR-518d-5p | CUCUAGAGGGAAGC$\underline{\text{A}}$CUUUCUG | CUCUAGAGGGAAGC$\underline{\text{G}}$CUUUCUG | ...(((((((.....))))))) | ...(((((((.....))))))) | 0.084 | $3.2 \times 10^{-4}$ |
| hsa-miR-548au-5p | has-miR-548ar-5p | AAAAGUAAUUGC$\underline{\text{A}}$GUUUUUGC | AAAAGUAAUUGC$\underline{\text{G}}$GUUUUUGC | ..................... | ..........(((.....))) | 0.26 | -0.15 |
| hsa-miR-548au-5p | has-miR-548ay-5p | AAAAGUAAUUG$\underline{\text{U}}$GGUUUUUGC | AAAAGUAAUUG$\underline{\text{C}}$GGUUUUUGC | ..................... | ..................... | 0.31 | 0.18 |
| hsa-miR-519a-5p | has-miR-526a-5p, hsa-miR-520c-5p, hsa-miR-518d-5p | CUCUAGAGGGAAGC$\underline{\text{A}}$CUUUCUG | CUCUAGAGGGAAGC$\underline{\text{G}}$CUUUCUG | ...(((((((.....))))))) | ...(((((((.....))))))) | 0.084 | $3.2 \times 10^{-4}$ |

Table 16: Results for various criteria applied to the entries from miRNASNP-v3 that concern hsa-miR-4537. The table contains areas under the ROC curves ($A_{\mathrm{ROC}}$), as well as $p$-values based on the two-sided Mann-Whitney test. $p$-values of less than 0.05, which indicate criteria that perform significantly better than the random predictor (with $A_{\mathrm{ROC}} = 0.5$), in **bold**.

| Criterion | $A_{\mathrm{ROC}}$ | $p$-value |
|---|---|---|
| $\Delta p_{\mathrm{unfolded}}$ | 0.746 | $\mathbf{5.17 \times 10^{-3}}$ |
| $\Delta p_{\mathrm{unfolded\ seed}}$ | 0.509 | $9.27 \times 10^{-1}$ |
| $\Delta S^{(\mathrm{i})}$ | 0.473 | $7.70 \times 10^{-1}$ |
| $\langle S_{\mathrm{mut}} \rangle$ | 0.744 | $\mathbf{5.57 \times 10^{-3}}$ |
| $\langle \Delta S \rangle$, all cases | 0.744 | $\mathbf{5.57 \times 10^{-3}}$ |
| $\langle d_{\mathrm{Hamming}} \rangle L^{-1}$ | 0.289 | $\mathbf{1.76 \times 10^{-2}}$ |

### 6 Case studies

#### 6.1 hsa-miR-4537

We take the example of one of hsa-miR-4537, one of the individual miRNAs that exhibit a significant effect of secondary structure after adjusting for multiple hypothesis testing. The database contains a total of 20 entries for disease-associated mutations affecting this miRNA, out of which there are 16 unique point mutants with an unaffected seed region.

In Figure 17 we demostrate that all applicable criteria except for $\Delta p_{\mathrm{unfolded}}$ and $\Delta p_{\mathrm{unfolded\ seed}}$ perform significantly better than random for this individual microRNA. We do not show the results for the two criteria based on $\langle d_{\mathrm{Hamming}} \rangle$ separately because the percentile ranking is fully equivalent to the one by the normalized Hamming distance. This likely indicates that, while the effect of secondary structure is by no means the only factor that determines the association of miRNAs with disease, it plays a significant role in some cases. Moreover, hsa-mir-4537 is a tumour-suppressor miRNA relevant to gastric cancer [72], and, as one would expect if the secondary structure affected activity, the mutants that change the folding most tend to be associated with disease, leading to $A_{\mathrm{ROC}} > 0.5$ for $\Delta p_{\mathrm{unfolded}}$ and $A_{\mathrm{ROC}} < 0.5$ for $\langle d_{\mathrm{Hamming}} \rangle L^{-1}$. As one would expect if the secondary structure affected activity, the mutants that change the folding most tend to be associated with disease, leading to $A_{\mathrm{ROC}} > 0.5$ for $\Delta p_{\mathrm{unfolded}}$ and $A_{\mathrm{ROC}} < 0.5$ for $\langle d_{\mathrm{Hamming}} \rangle L^{-1}$. We provide additional data on this miRNA in Figure 18, which contains the ROC curves underlying Figure 17, and Table 16, which gives the values of $A_{\mathrm{ROC}}$ for the various criteria.

#### 6.2 hsa-miR-485-5p

We also present the results of applying the various criteria to data for hsa-miR-485-5p, for which the $p$-value that quantifies the significance of the result for $\langle d_{\mathrm{Hamming}} \rangle / L$ is the lowest ($p = 5.45 \times 10^{-3}$). The miRNASNP-v3 database contains a total of 6 entries for disease-associated mutations affecting this miRNA, out of which there are 4 unique point mutants with an unaffected seed region.

As Figure 19 illustrates, all of our criteria except for $\Delta p_{\mathrm{unfolded}}$ and $\Delta p_{\mathrm{unfolded\ seed}}$ perform significantly better than random for this individual microRNA. This provides further support for the hypothesis that secondary structure impacts the function of some miRNAs. This miRNA is known to act as an inhibitor of breast cancer progression [49]; despite that, it is the mutants with secondary structure close to that of the WT that tend to be associated with disease, as indicated by the $A_{\mathrm{ROC}}$ value for $\langle d_{\mathrm{Hamming}} \rangle / L$ in the figure ($A_{\mathrm{ROC}} = 0.926$). We provide additional data on this miRNA in Figure 20, which contains the ROC curves underlying Figure 19, and Table 17, which gives the relevant areas under the ROC curves.

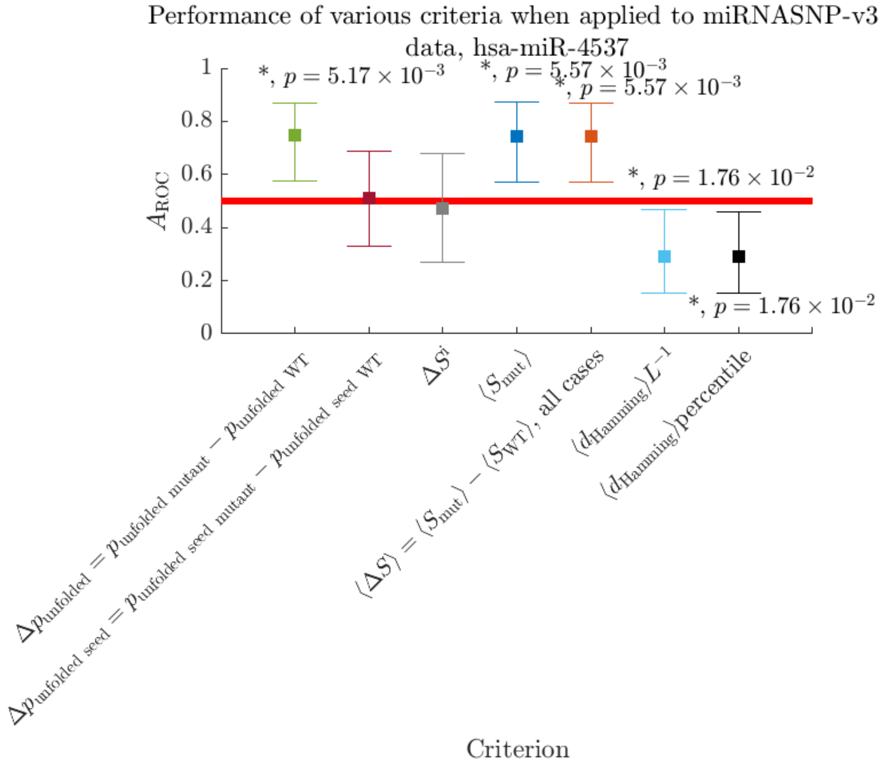

Figure 17: **Multiple independent SS-based criteria predict association of miRNA mutations with disease better than random for hsa-miR-4537.** Comparison of the performance of various criteria in terms of predicting disease-associated mutations when applied to mutations concerning hsa-miR-4537 from miRNASNP-v3. hsa-miR-4537 is known to play a tumour-suppressing role in gastric cancer [72]. Squares mark $A_{\text{ROC}}$ values, error bars show 95% confidence intervals calculated with the bootstrapping method [207], and Mann-Whitney $p$-values are indicated wherever $p < 0.05$. The red horizontal line indicates the area under the curve for the random criterion, $A_{\text{ROC}} = 0.5$. The only criteria that do not perform significantly better than the random one for this dataset are $\Delta p_{\text{unfolded seed}}$, which measures the change in probability that the miRNA seed region is fully unfolded and $\delta S^{\text{i}}$, which measures the change in the positional entropy of the mutated site. The comparatively large values of $A_{\text{ROC}}$ suggest that miRNA folding plays a role in disease and therefore miRNA activity.

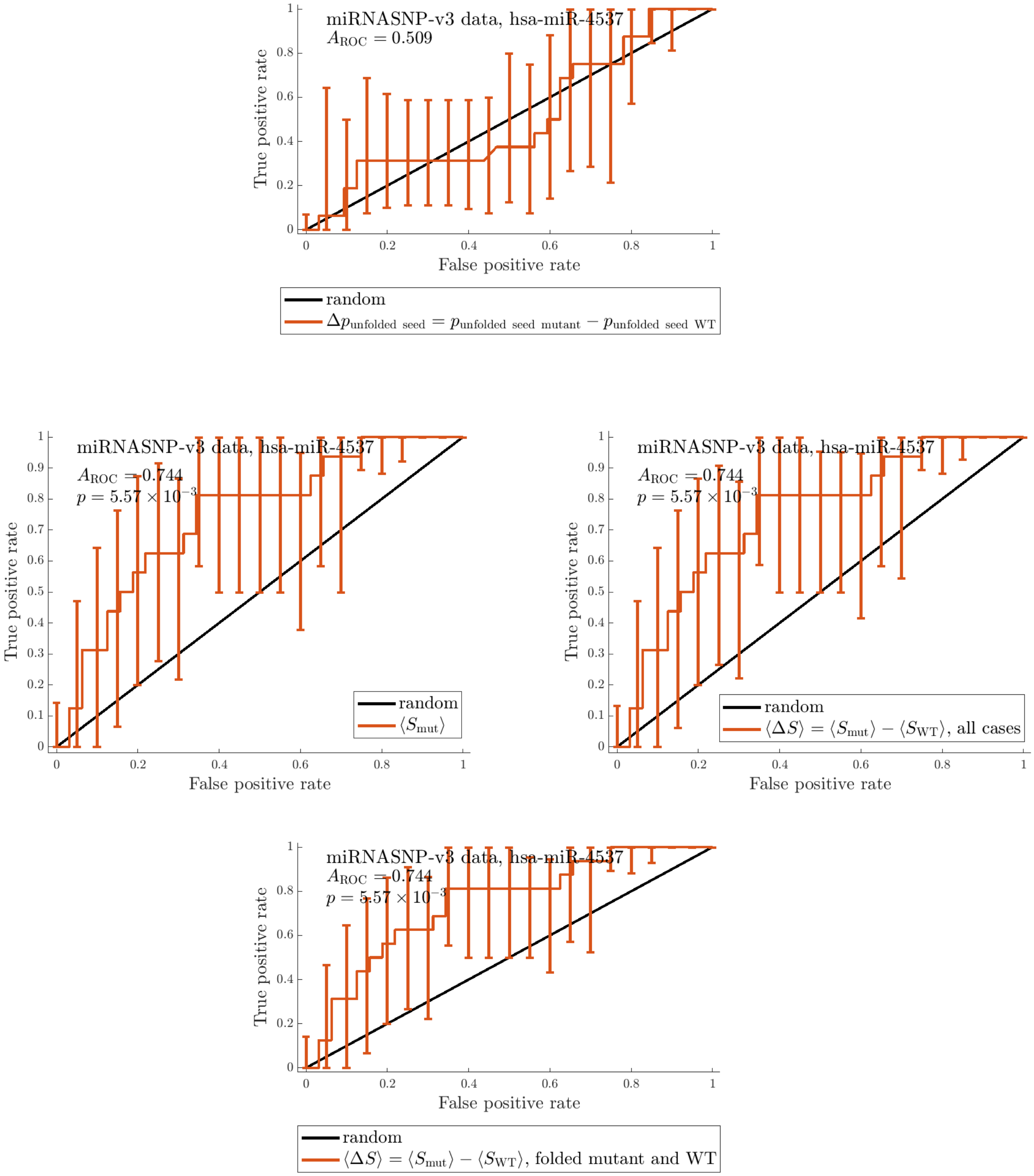

Figure 18: ROC curves characterizing the performance of all criteria discussed above for mutations affecting hsa-miR-4537 from miRNASNP-v3.

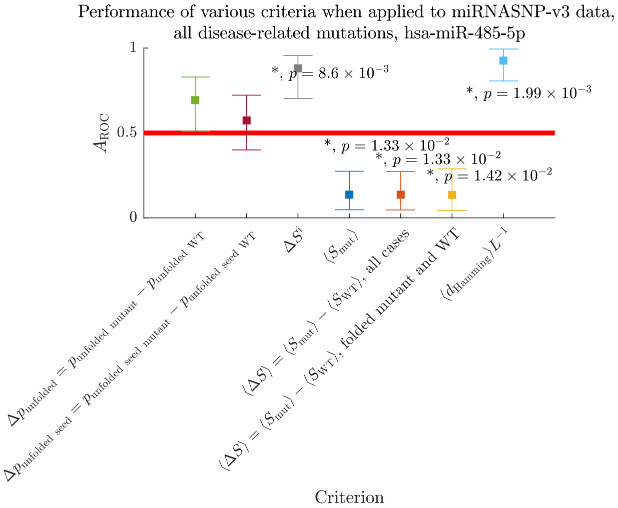

Figure 19: **Multiple independent SS-based criteria predict association of miRNA mutations with disease better than random for hsa-miR-485-5p.** Comparison of the performance of various criteria in terms of predicting disease-associated mutations when applied to mutations concerning hsa-miR-485-5p from miRNASNP-v3. hsa-miR-485-5p is associated with various types of cancer, including lung cancer, breast cancer and others [42–54] and cerebral ischemia [55]. Squares mark $A_{\mathrm{ROC}}$ values, error bars show 95% confidence intervals, and Mann-Whitney $p$-values are indicated wherever $p < 0.05$. The red horizontal lines indicate the area under the curve for the random criterion, $A_{\mathrm{ROC}} = 0.5$. The only criteria that do not perform significantly better than the random one for this dataset are those based on the probability of folded states. The comparatively large values of $A_{\mathrm{ROC}}$ suggest that miRNA folding plays a role in disease and therefore miRNA activity.

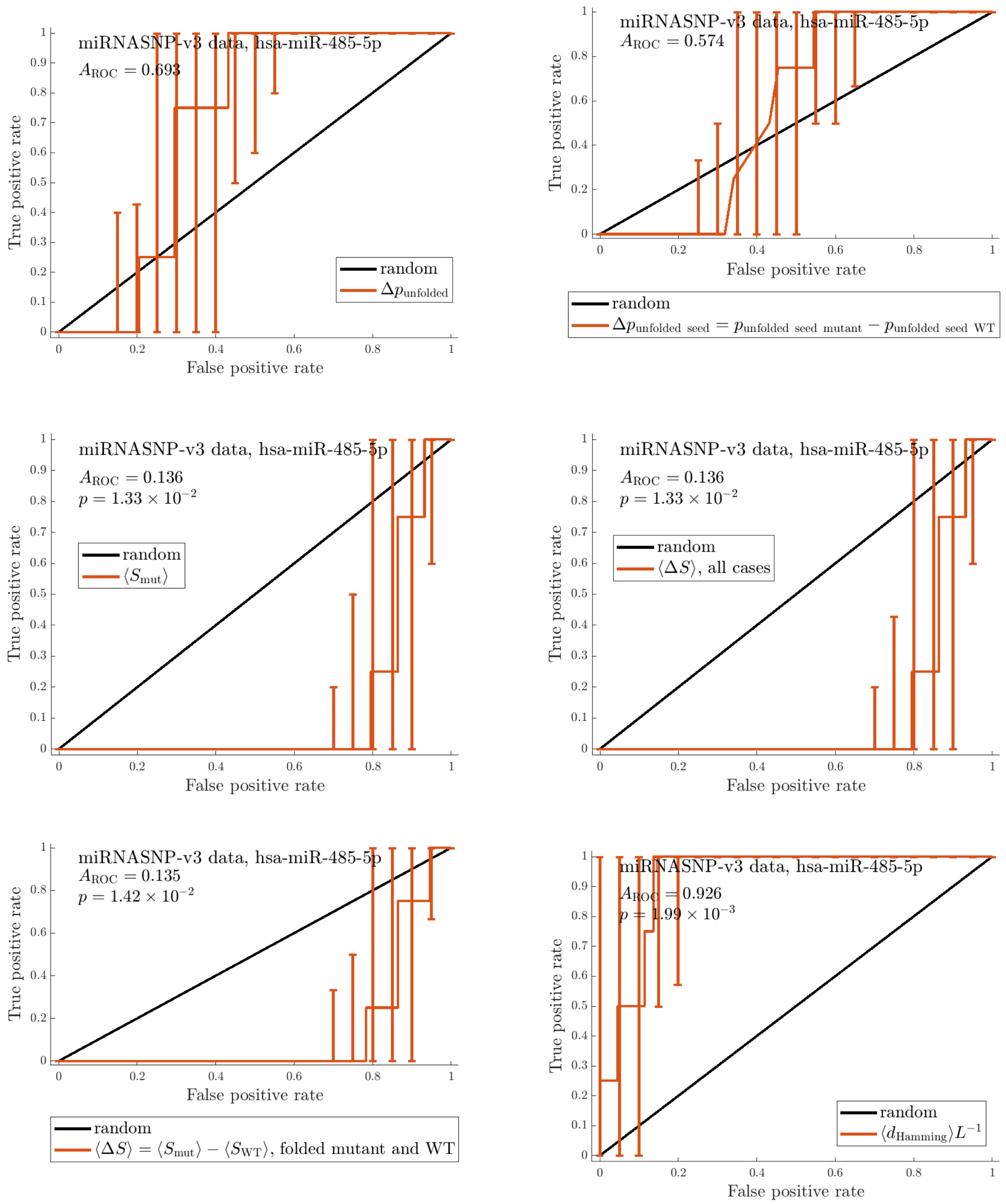

Figure 20: ROC curves characterizing the performance of all criteria discussed above for mutations affecting hsa-miR-485-5p from miRNASNP-v3.

Table 17: Results for various criteria applied to the entries from miRNASNP-v3 that concern hsa-miR-485-5p. The table contains areas under the ROC curves ($A_{\mathrm{ROC}}$), and $p$-values based on the two-sided Mann-Whitney test. $p$-values of less than 0.05, which indicate criteria that perform significantly better than the random predictor (with $A_{\mathrm{ROC}} = 0.5$), in **bold**.

| Criterion | $A_{\mathrm{ROC}}$ | $p$-value |
|---|---|---|
| $\Delta p_{\mathrm{unfolded}}$ | 0.693 | $2.21 \times 10^{-1}$ |
| $\Delta p_{\mathrm{unfolded\ seed}}$ | 0.574 | $6.47 \times 10^{-1}$ |
| $\Delta S^{(\mathrm{i})}$ | 0.881 | $\mathbf{8.60 \times 10^{-3}}$ |
| $\langle S_{\mathrm{mut}} \rangle$ | 0.136 | $\mathbf{1.33 \times 10^{-2}}$ |
| $\langle \Delta S \rangle$, all cases | 0.136 | $\mathbf{1.33 \times 10^{-2}}$ |
| $\langle d_{\mathrm{Hamming}} \rangle L^{-1}$ | 0.926 | $\mathbf{1.99 \times 10^{-3}}$ |

### 7 Additional information on miRNAs with $q < 0.05$

When we apply the strictest level of filtering, $N_{\mathrm{mut\ seed}} \geq 3$ and $N_{\mathrm{mut\ non-seed}} \geq 3$, to the dataset with mutations associated with any traits and diseases, we find one miRNA with a $q$-value that passes our significance threshold for the criterion based on the mutant-WT SS Hamming distance $\langle d_{\mathrm{Hamming}} \rangle / L$ criterion - $q = 3.72 \times 10^{-2}$ for **hsa-miR-4477b**.

Additionally, we calculate the Benjamini-Hochberg $q$-values for the combined $p$-values calculated via the Fisher method [208] for $\Delta p_{\mathrm{folded}}$ and $\langle d_{\mathrm{Hamming}} \rangle / L$ of the individual miRNAs represented in SomamiR and miRNASNP-v3. When we analyse the mutations from miRNASNP-v3 that pertain to all diseases and traits with applied filtering by $N_{\mathrm{mut\ seed}}$ and $N_{\mathrm{mut\ non-seed}}$, we find $q$-values are below 0.05 for several miRNAs. At $N_{\mathrm{mut\ seed}} \geq 1$ and $N_{\mathrm{mut\ non-seed}} \geq 1$, $q = 2.67 \times 10^{-2}$ for **hsa-miR-4641** and **hsa-miR-1269b**. For the smaller subset with $N_{\mathrm{mut\ seed}} \geq 2$ and $N_{\mathrm{mut\ non-seed}} \geq 2$, $q = 1.86 \times 10^{-2}$ for **hsa-miR-4537** and **hsa-miR-4477b**, $q = 1.10 \times 10^{-2}$ for hsa-miR-1269b, and the $q$-value is further reduced to $q = 1.31 \times 10^{-2}$ for **hsa-miR-4537** and **hsa-miR-4477b** based on the set with $N_{\mathrm{mut\ seed}} \geq 3$ and $N_{\mathrm{mut\ non-seed}} \geq 3$.

### 8 Code and data

We provide an archive with the code used for generating the results discussed in the paper, as well as the main results themselves in the form of tables and figures.
